# Canalized gene regulatory networks stabilize floral polymorphism and enable modular transgressive expression

**DOI:** 10.64898/2026.04.16.719044

**Authors:** Santosh K Rana, Hum K Rana, Jacob B Landis, Juntong Chen, Tao Deng, Hang Sun

## Abstract

- Floral polymorphisms frequently persist across heterogeneous environments despite ongoing gene flow, yet the regulatory mechanisms maintaining discrete phenotypes remain unclear. We tested whether alternative flower-colour morphs in *Stellera chamaejasme* L. are maintained by canalized gene regulatory architectures that stabilize expression around morph-specific optima.
- We used a pan-transcriptomic and eco-evolutionary framework integrating genome-wide gene expression profiling, co-expression network analysis, functional enrichment, ortholog-based phylogenomics, and variance-based modeling of regulatory canalization and transgressive expression to quantify regulatory variation across morphs.
- Transcriptomic variation was structured primarily by morph identity rather than geography, indicating consistent morph-associated regulatory programs. Parental morphs showed reduced within-morph variance in gene co-expression modules, consistent with strong regulatory constraint at the network level. In contrast, a naturally occurring mosaic morph exhibited extensive non-additive and predominantly transgressive expression, with most genes falling outside the parental range. This transgressive signal was modular, with most networks remaining stable while a subset showed elevated variance and disrupted inheritance. Functional analyses further reveal that floral pigmentation is embedded within broader metabolic and stress-response pathways, linking color polymorphism to coordinated physiological states and ecological differentiation.

## Introduction

Understanding how discrete phenotypic polymorphisms persist across populations despite ongoing gene flow remains a central challenge in evolutionary biology and ecological genomics (Gray & McKinnon, 2007; Kitano *et al*., 2025). In theory, recombination and migration should homogenize phenotypic variation, yet many natural systems maintain distinct and stable morphs across heterogeneous landscapes. Flower-color polymorphisms provide particularly powerful models for addressing this ambiguity because they are visually discrete, ecologically consequential, and repeatedly associated with pollinator behavior, abiotic stress, and reproductive isolation across diverse plant lineages (Schemske & Bradshaw, 1999; Hopkins & Rausher, 2012; Wessinger *et al*., 2014). Although phenotypic differentiation among floral morphs is well documented, the regulatory mechanisms that allow alternative phenotypes to persist across spatially structured populations remain poorly understood.

A key unresolved question is whether such polymorphisms are maintained primarily by selection acting on individual loci or by stabilization of higher-order gene regulatory architectures. Eco-evolutionary theory predicts that when alternative phenotypes are adaptive, stabilizing or phenotype-associated selection may act on gene regulatory networks, producing canalized transcriptional states with reduced variance around morph-specific optima (Waddington, 1942; Gibson & Wagner, 2000; Masel & Siegal, 2009; Paaby & Rockman, 2014). Under this framework, divergence among morphs should be reflected not only in differences in mean expression, but also in reduced within-morph variance and recurrent regulatory configurations across populations despite gene flow (Yeaman & Whitlock, 2011; Tigano & Friesen, 2016; Ravinet *et al*., 2017). Because complex traits are controlled by coordinated interactions among many genes, such stability is expected to emerge at the level of gene regulatory networks and co-expression modules rather than individual loci (Wagner, 2011; Boyle *et al*., 2017; Payne & Wagner, 2019; Kadelka, 2026).

A major challenge, however, is distinguishing phenotype-associated regulatory structure from patterns generated by neutral spatial processes. Geographic distance, demographic history, and spatially autocorrelated environments can generate transcriptional divergence through drift or plasticity alone, potentially leading to false inference of adaptive divergence if spatial structure is not explicitly modeled (Meirmans, 2012; Wang & Bradburd, 2014; Rellstab *et al*., 2015; Forester *et al*., 2018; Dauphin *et al*., 2023). Robust inference therefore requires analytical frameworks that explicitly partition transcriptomic variation into phenotype-associated (selection-consistent) and geography-associated (drift-consistent) components (Peres-Neto *et al*., 2006; Capblancq *et al*., 2020; Li *et al*., 2025). If regulatory canalization underlies the persistence of floral morphs, transcriptomic similarity should align more strongly with morph identity than with geographic proximity, producing a pattern increasingly described as isolation by morph rather than isolation by distance. Perturbations of stabilized regulatory systems provide a complementary approach for testing the limits of canalization (Horta-Lacueva *et al*., 2023; Runemark *et al*., 2025). When divergent regulatory backgrounds are combined or reorganized, buffering may be relaxed, exposing epistatic interactions and generating non-additive or transgressive expression (Dobzhansky, 1982; Rieseberg *et al*., 1999; McManus *et al*., 2010; Runemark *et al*., 2025). Importantly, such responses are often modular, with core regulatory components remaining stable while specific subnetworks become destabilized (Gibson & Wagner, 2000; Wagner, 2008; Félix & Barkoulas, 2015; Kadelka & Murrugarra, 2024).

Floral pigmentation provides an ideal system for testing these ideas because it reflects integrated metabolic and regulatory programs rather than the action of a single locus. Anthocyanin and carotenoid biosynthesis pathways are tightly linked with primary metabolism, redox balance, stress responses, and developmental regulation, making floral color a pleiotropic trait embedded within broader physiological networks (Rausher, 2008; Albert *et al*., 2014; Smith, 2016). Consequently, canalization of pigmentation is expected to arise from stabilization of these broader regulatory systems, whereas disruption of such networks may produce expression instability and phenotypes extending beyond the parental range (Smith & Rausher, 2011; Wessinger & Rausher, 2015; Chen *et al*., 2025).

Here, we investigate these dynamics in *Stellera chamaejasme* L., a widespread alpine perennial of the Qinghai–Tibetan Plateau and adjacent montane regions that exhibits four stable parental flower-color morphs of pure pink (PP), red–white (RW), yellow–white (YW), and pure yellow (YY), distributed across broad geographic and elevational gradients (Rana *et al*., 2024; Sun *et al*., 2024). We additionally identify a naturally occurring mosaic morph (RWh) in zones of sympatry between PP and RW populations, providing a natural perturbation of regulatory architecture for testing whether gene expression follows additive expectations or exhibits transgressive reorganization. Using a pan-transcriptomic framework integrating RNA-seq data across morphs, tissues, replicated populations, and geographic regions, we test three predictions: (i) transcriptomic divergence is structured more strongly by morph identity than by geography; (ii) parental morphs exhibit reduced within-population variance consistent with transcriptional canalization; and (iii) the RWh morph reveals the limits of regulatory stability through additive, dominant, or transgressive expression relative to parental morphs. Together, this framework allows us to evaluate how canalized regulatory networks contribute to the persistence of floral polymorphism and how its disruption generates transgressive regulatory states.

## Materials and Methods

### Study system for eco-evolutionary framework

*Stellera chamaejasme* is a widespread alpine perennial of the Qinghai–Tibetan Plateau and adjacent montane regions. This species exhibits discrete flower-color polymorphism, including pure pink (PP), red–white (RW), yellow–white (YW), and pure yellow (YY) (Rana *et al*., 2024). The persistence of these morphs, despite potential gene flow, makes this system well suited for testing how stabilizing selection structures regulatory architecture across populations. During field sampling, we identified a naturally occurring red–white mosaic morph (RWh) in zones of sympatry between PP and RW populations. These individuals frequently produced inflorescences bearing both RW-like and PP-like floral elements, suggesting a probable hybrid origin from these two parental morphs. One hypothesis is that RWh does not occupy an intermediate position in transcriptomic expression space but instead exhibits extensive regulatory divergence, consistent with a strongly transgressive expression state. Yellow-based morphs (YW and YY) were not observed in association with these mixed inflorescences and were therefore treated as non-parental reference morphs in subsequent analyses. This system, therefore, provides a natural eco-evolutionary framework to test whether floral morphs correspond to stabilized transcriptional states, whether such states are canalized within populations, and how hybridization and transgressive regulatory states may perturb stabilized regulatory architectures (Fig. 1).

**Figure 1.**
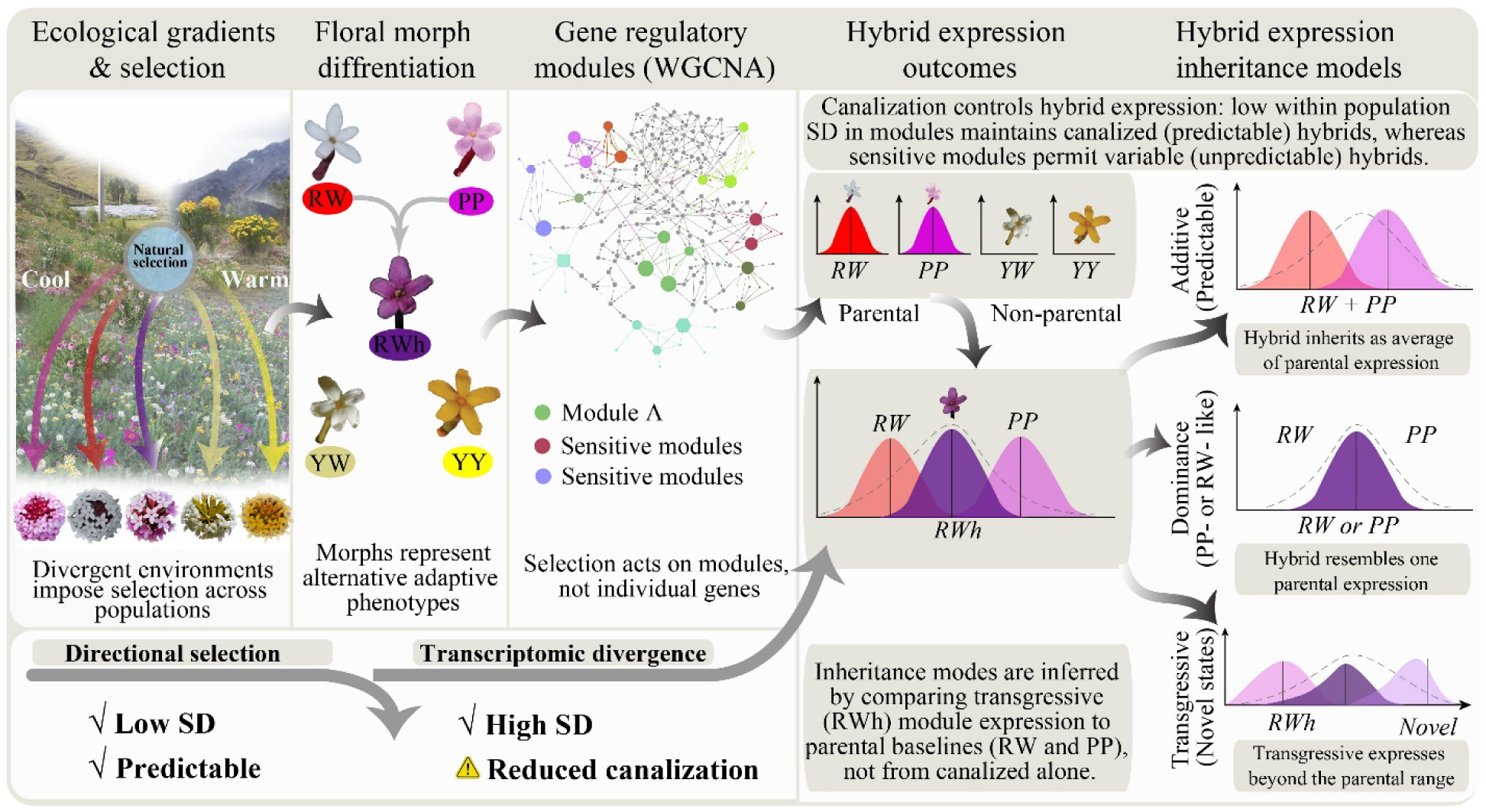
Eco-evolutionary framework linking floral polymorphism, regulatory canalization, and hybrid predictability. Conceptual framework illustrating how ecological divergence is associated with transcriptional regulatory structure and expression outcomes in a mosaic morph (RWh). Floral morphs (PP, RW, YW, YY) represent distinct phenotypic states hypothesized to correspond to canalized gene regulatory networks, where co-expression modules exhibit reduced within-population variance. A rare mosaic morph (RWh), occurring in zones of sympatry between PP and RW, represents a perturbation of these regulatory states. Expression in RWh may be additive or parent-biased in strongly canalized modules, whereas less constrained modules may exhibit non-additive or transgressive expression. Thus, most modules remain stable, while a subset becomes destabilized, generating transgressive regulatory states. Arrows indicate the progression from ecological divergence to regulatory architecture and expression outcomes.

### Field sampling and phenotypic characterization

For each floral morph (PP, RW, YW, YY), three geographically distinct populations (at least 1 km far apart) were sampled across Southwest China along elevational gradients (2,676–4,210 m a.s.l; Fig. 2A; Supplementary Table S1). Within each population, flower tissue from three individuals was collected to provide biological replication, with sampled plants spaced at least 100 m apart to minimize the likelihood of sampling closely related or clonally derived individuals. The RWh morph was sampled from two sympatric populations, reflecting its rarity. Floral tissue was collected at comparable developmental stages to minimize ontogenetic effects on gene expression and was immediately flash-frozen in liquid nitrogen to ensure RNA integrity. Leaf, stem, and root tissues were additionally collected to support transcriptome assembly. Geographic coordinates were recorded for all populations, and voucher specimens were deposited at the Herbarium of the Kunming Institute of Botany (KUN).

**Figure 2.**
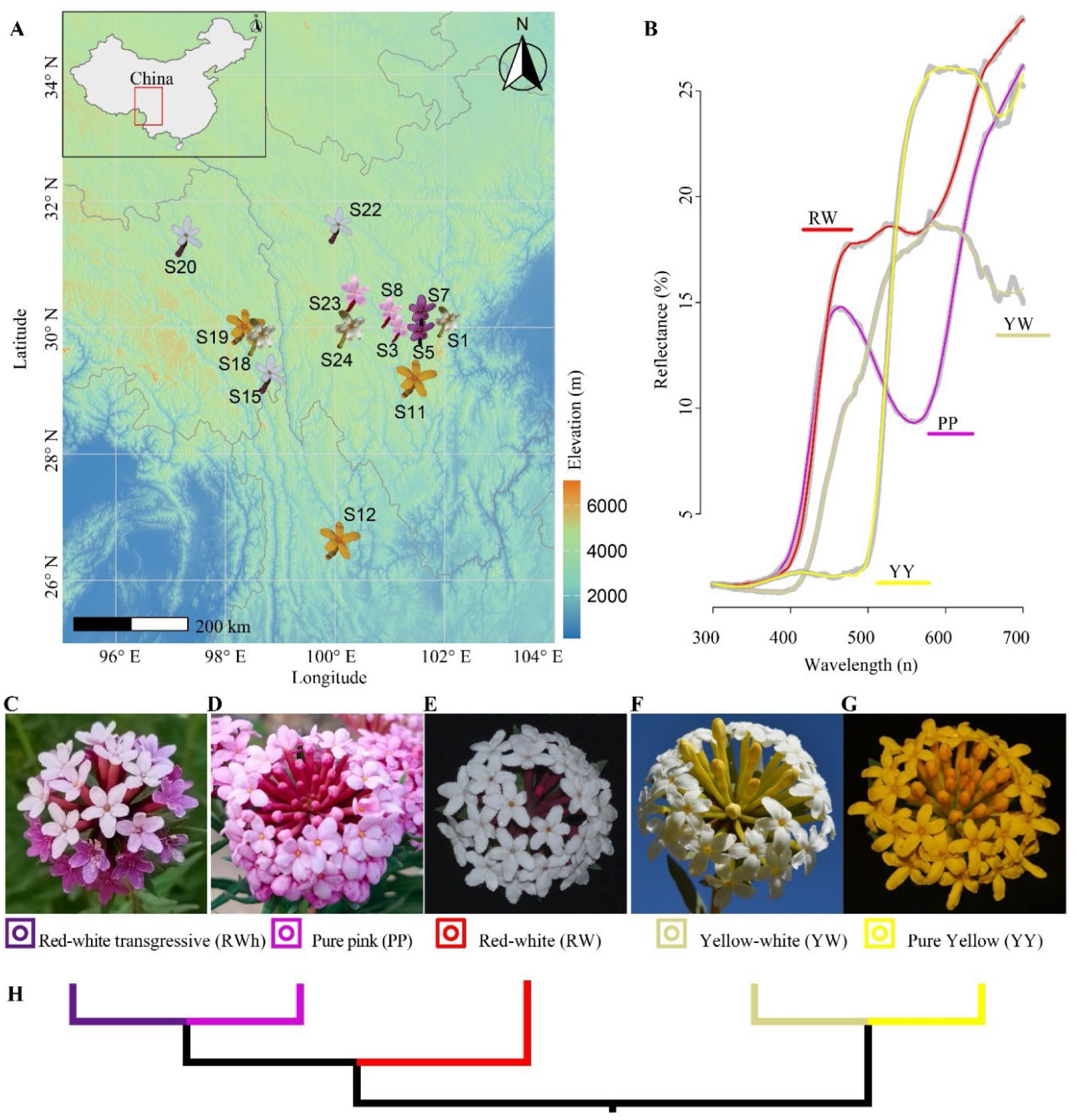
Geographic distribution, phenotypic differentiation, and phylogenetic relationships of floral morphs in *Stellera chamaejasme*. **(A)** Geographic distribution of sampled populations across southwestern China. Sampling sites are indicated by flower icons colored by dominant morph; background shading represents elevation (m a.s.l.). Inset shows the study region within China. **(B)** Mean floral reflectance spectra (300–700 nm) for four morphs (PP, RW, YW, YY), showing distinct spectral profiles across the visible range. (**C**–**G**) Representative inflorescences of the five morphs: (**C**) pure pink (PP), (**D**) red–white (RW), (**E**) red–white mosaic (RWh), (**F**) yellow–white (YW), and (**G**) pure yellow (YY). Symbols correspond to morph designations used throughout. (**H**) Morph-level phylogeny inferred from genome-wide single-copy orthologs (OrthoFinder), showing relationships among morphs. The yellow lineage (YW, YY) diverges first, followed by separation of RW from the pink lineage, with PP and RWh sharing the most recent common ancestry.

Floral spectral reflectance (300–700 nm) was measured for representative individuals using a fiber-optic spectrometer (USB2000+, Ocean Optics, Dunedin, FL, USA) equipped with a deuterium–halogen UV–VIS light source (DH-2000, Ocean Optics), with measurements calibrated against a pressed barium sulfate (BaSO₄) pellet serving as a diffuse white reflectance standard. Spectral data were processed in the R-package *pavo2* v2.7.0 (Maia *et al*., 2019), providing quantitative phenotypic validation of discrete floral morphs.

### RNA sequencing, transcriptome assembly, and expression quantification

Total RNA was extracted and purified from the 1 mg of flower, leaf, stem, and root tissues using the Spectrum^TM^ Plant Total RNA Kit (STRN250, Sigma-Aldrich). RNA concentration was assessed using the Qubit RNA Assay Kit with a Qubit 2.0 Fluorometer (Life Technologies, CA, USA) and purity was assessed with a NanoDrop2000 spectrophotometer (Thermo Fisher Scientific, Waltham, MA, USA). Strand-specific cDNA libraries were prepared using the NEBNext Ultra^TM^ RNA/DNA Library Prep Kit (New England Biolabs, USA) and sequenced on an Illumina HiSeq^TM^ 2500 platform using paired-end (2 × 150 bp), generating raw reads of 19.58 to 33.66 million reads per library. In total, 160 libraries were generated (144 parental; 16 transgressive morphotype) (Table S2). Library preparation and sequencing were performed by Novogene Bioinformatics Technology Co., Ltd. (Beijing, China).

A unified pan-transcriptome was constructed from all morphs and tissues. Raw paired-end reads were quality filtered using *Trimmomatic* v0.39 (Bolger *et al*., 2014) to remove adapter sequences and low-quality bases using sliding-window trimming (4 bp window, minimum Phred score of 20), with reads shorter than 50 bp discarded. Following filtering, clean reads from flower, leaf, stem, and root tissues were concatenated for each individual to generate a single paired-end dataset. These individual-level read sets were subsequently pooled across all samples for *de novo* assembly. The pan-transcriptome was assembled using *Trinity* v2.13.2 (Grabherr *et al*., 2011; Haas *et al*., 2013) with strand-specific parameters. In addition to the global assembly, tissue-specific (flower, leaf, stem, and root) and morph-specific (PP, RW, RWh, YW, and YY) assemblies were generated by pooling reads within tissue or morph categories to evaluate assembly consistency and transcript representation across biological groups. Redundant transcripts were removed using *CD-HIT-EST* v4.8.1 (Fu *et al*., 2012) using a 95% identity threshold to generate a non-redundant reference assembly. Assembly contiguity was evaluated using Trinity’s expression-weighted ExN50 statistics, and completeness was assessed with *BUSCO* v5.4.7 (Simão *et al*., 2015) in transcriptome mode against the embryophyta_odb10 lineage dataset, reporting the proportions of complete (single-copy/duplicated), fragmented, and missing conserved plant orthologs.

Coding regions were predicted from the non-redundant transcriptome using *TransDecoder* v5.7.1(Haas *et al*., 2013). Functional annotation was performed by similarity searches against the UniProt/SwissProt protein database using BLASTX (Camacho *et al*., 2009) with an E-value threshold of 1 × 10⁻⁵. Predicted protein sequences were further annotated using *eggNOG-mapper* v2 (Cantalapiedra *et al*., 2021) to assign Gene Ontology (GO) terms, KEGG pathways, and orthologous functional categories based on precomputed orthologous groups.

A Salmon index was constructed from the non-redundant pan-transcriptome assembly. Transcript abundance was estimated using *Salmon* v1.8.0 (Patro *et al*., 2017) in quasi-mapping (mapping-based) mode, with reads mapped directly to the transcriptome without prior alignment. Transcript-level abundance estimates were summarized as transcripts per million (TPM) and raw estimated counts. Gene-level counts were generated using Trinity gene–transcript mappings. For differential expression analyses, raw gene-level counts were normalized using the median-of-ratios method in the R-package *DESeq2* v1.42 (Love *et al*., 2014). Differential expression was assessed using generalized linear models, with multiple-testing correction applied using the Benjamini–Hochberg false discovery rate procedure. For co-expression and eco-evolutionary analyses, expression data were normalized using trimmed mean of M-values (TMM) normalization in the R-package *edgeR* v3.44 (Robinson *et al*., 2010) and log_2_-transformed.

All expression processing steps were performed using reproducible workflows implemented in R v4.3, with key R-packages including *DESeq2* v1.42*, edgeR* v3.44*, limma* v3.58, and *tidyverse* v2.0. Normalization choices were tailored to the analytical objective of inferring gene-level inference for differential expression and module-level inference for regulatory and eco-evolutionary analyses, ensuring methodological consistency while maximizing biological interpretability.

### Early test of transgressiveness in floral expression and sequence space

Field observations showed that the RWh exhibits a mosaic floral phenotype combining traits of the sympatric morphs PP and RW, with several trait values extending beyond the parental range. Based on this pattern, RWh was presumed to represent a transgressive hybrid, which we evaluated using integrated orthologous phylogenomic and expression-based analyses. To assess the evolutionary placement of the RWh morphotype relative to other morphs, a morph-level phylogenetic analysis was conducted based on genome-wide orthologs. Non-redundant transcript assemblies generated after CD-HIT-EST clustering were translated into predicted coding sequences using *TransDecoder* v5.7.1 (Haas *et al*., 2013). The *OrthoFinder* v2.5 (Emms & Kelly, 2019) was used to infer orthogroups and reconstruct a morph-level species tree. Maximum-likelihood gene trees were estimated for each orthogroup and combined using the STAG algorithm to produce a genome-wide consensus topology, which was rooted using STRIDE (Emms & Kelly, 2019).

Immediately after constructing the filtered gene-level expression matrix, we evaluated whether the RWh morph exhibits intermediate expression consistent with simple parental inheritance or a transgressive regulatory state relative to the two morphs observed in sympatry (PP and RW) using a hybrid index (HI). Gene-level expression values were matched to morph identity, and genes with low expression (mean ≤ 1 across samples) were removed. For each gene, *g*, mean expression was calculated for each morph, and a hybrid index was computed along the parental PP–RW axis as:

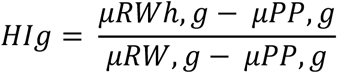

Where, *μ*_*m*,_*_g_* is the mean expression of gene *g* in morph m. This formula follows hybrid index approaches used to quantify genomic or expression inheritance along parental gradients (Landry *et al*., 2007; McManus *et al*., 2010). Genes with *μ*_RW,_*_g_* = *μ_PP_*,*_g_* were excluded for that gene because the denominator was zero. Values between 0 and 1 indicate expression within the parental range, whereas values <0 or >1 indicate transgressive expression outside the parental range. To account for differences at the gene expression scale, the analysis was repeated after gene-wise standardization (z-scoring), allowing detection of transgressiveness in relative expression profiles rather than absolute abundance. Genes were classified into inheritance categories based on hybrid index thresholds: PP-like (0 ≤ *HI* ≤ 0.25), intermediate (0.25 < *HI* < 0.75), RW-like (0.75 ≤ *HI* ≤ 1), transgressive below PP (*HI* < 0), and transgressive above RW (*HI* > 1). Genome-wide distributions of hybrid index values were visualized using histograms, and category proportions were calculated to quantify the relative contribution of additive, dominant, and transgressive regulatory patterns.

### Transcriptomic analyses and regulatory architecture

Following the early assessment of genome-wide expression inheritance and transgressiveness, we conducted comprehensive analyses to characterize morph-associated regulatory divergence and the transcriptional architecture of the RWh lineage. Gene-level differential expression among floral morphs were assessed using the R-package *DESeq2* v1.42 (Love *et al*., 2014), with morph identity specified as the focal factor and population included as a blocking factor, where appropriate. Count data were modeled using a negative binomial framework, and significance was evaluated using Wald tests with Benjamini–Hochberg false discovery rate (FDR) correction. Differentially expressed genes provided an initial measure of morph-associated regulatory divergence and informed downstream functional and network analyses.

To identify coordinated regulatory programs, weighted gene co-expression networks were constructed using the R-package *WGCNA* v1.72-5 (Langfelder & Horvath, 2008) using TMM-normalized, log_2_-transformed floral expression data. An unsigned network was inferred using Pearson correlations, with soft-thresholding parameters chosen to approximate a scale-free topology. Co-expression modules were identified based on topological overlap and defined using dynamic tree cutting. Module eigengenes (first principal component of module expression) were used as low-dimensional summaries of regulatory activity. Module eigengenes formed the basis for all downstream eco-evolutionary analyses.

Functional interpretation of transcriptional divergence was performed at the gene level using the R-package *clusterProfiler* v4.0 (Yu *et al*., 2012), testing Gene Ontology (GO) and KEGG pathway enrichment (Kanehisa, 2000) for differentially expressed genes and WGCNA modules. Enrichment significance was assessed using Benjamini–Hochberg FDR correction. Analyses focused on pathways associated with floral pigmentation, secondary metabolism, plastid function, and regulatory processes to facilitate biological interpretation of morph-associated and RWh-specific regulatory patterns.

### Partitioning morph-associated divergence versus geographic structure

To evaluate whether transcriptomic variation aligns more strongly with floral phenotype than with spatial structure, module eigengenes were averaged within populations to generate population-level regulatory profiles. Euclidean distance matrices were analyzed using permutational multivariate analysis of variance (PERMANOVA) implemented in the *adonis2* function in the R-package *vegan* v2.6-4 (Oksanen *et al*., 2019) using 999 permutations, with morph identity as the explanatory factor. Spatial structure was modeled using spatial eigenvectors derived from principal coordinates of neighbor matrices (PCNM) calculated from geographic distances among populations (Borcard *et al*., 2018).

Distance-based redundancy analysis (dbRDA) was then performed using the *capscale* function in the R-package *vegan* v2.6-4 (Oksanen *et al*., 2019), with morph identity as the explanatory factor and spatial eigenvectors included as conditional variables. Morph identity was included as the focal predictor, with spatial eigenvectors specified as conditional variables. Significance of constrained models and marginal effects was assessed using permutation tests.

Variance partitioning analysis was then used to quantify the unique and shared contributions of morph identity and geographic structure to transcriptomic divergence (Borcard *et al*., 1992; Peres-Neto *et al*., 2006). Adjusted *R^2^* values were reported to avoid inflation of explained variance. A greater proportion of variance uniquely attributable to morph identity relative to geography was interpreted as evidence that transcriptional divergence is structured primarily by phenotype-associated selection rather than neutral isolation by distance alone, consistent with recent eco-evolutionary genomic studies using multivariate expression data (Forester *et al*., 2018; Capblancq *et al*., 2020).

### Quantifying transcriptional canalization using within-population variance

Transcriptional canalization was quantified as reduced within-population variance of regulatory modules (Waddington, 1942; Gibson & Wagner, 2000). For each population and morph, within-population variance was calculated as the standard deviation (SD) of module eigengene expression across biological replicates. Module-level SDs were calculated separately for each population and morph using workflows implemented in the R-package *tidyverse* v2.0.0 (Wickham *et al*., 2019). Populations with only one replicate (n=1 in population S5; Table S1) were excluded from variance estimation but retained for population-mean analyses. Low within-population variance for a given module was interpreted as stronger regulatory constraint, consistent with stabilizing selection acting on gene regulatory networks (Rifkin *et al*., 2005; Landry *et al*., 2007; Masel & Siegal, 2009).

Distributions of module-level variance were compared among floral morphs using Wilcoxon rank-sum tests, with false discovery rate (FDR) correction applied using the Benjamini–Hochberg procedure (Benjamini & Hochberg, 1995). Reduced variance was interpreted as stronger regulatory constraint consistent with stabilizing selection (Landry *et al*., 2007; McManus *et al*., 2010). To quantify hybrid destabilization, we calculated a hybrid variance inflation index for each module as:

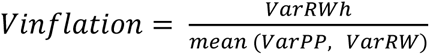

Values greater than one indicate reduced canalization relative to parental morphs, and potential regulatory incompatibility (Rieseberg *et al*., 1999; Mack & Nachman, 2017). Modules exhibiting elevated hybrid variance inflation were treated as candidates for regulatory destabilization and potential phenotypic novelty.

### Classifying hybrid inheritance at the regulatory module level

Regulatory inheritance was evaluated at the level of co-expression modules using population-mean module eigengenes for PP, RW, and RWh. Inheritance categories were defined as additive, dominant (PP-like or RW-like), or transgressive following established frameworks for RWh expression analysis (Landry *et al*., 2007; McManus *et al*., 2010). To minimize arbitrary thresholds, classification tolerance was defined using module-specific parental variance, calculated as half of the pooled parental standard deviation. This variance-informed approach aligns inheritance inference with the natural scale of regulatory variation for each module and reduces misclassification due to stochastic noise (Romero *et al*., 2012). Inheritance categories were summarized across modules and compared with variance inflation indices using Wilcoxon tests.

## Results

### Discrete floral morphs show stable geographic occurrence and functional differentiation

Field sampling across Southwest China identified four recurrent floral morphs of *Stellera chamaejasme*, pure pink (PP), red–white (RW), yellow–white (YW), pure yellow (YY), and a naturally occurring red–white mosaic morph (RWh) occurring in zones of sympatry between PP and RW populations (Fig. 2A; Supplementary Table S1). Each parental morph formed morphologically consistent populations across geographically separated sites spanning broad elevational gradients (2,676–4,210 m a.s.l.), indicating stable phenotypic recurrence across environmental and spatial contexts (Fig. 2A).

Floral reflectance measurements revealed clear spectral differentiation among parental morphs (PP, RW, YW, and YY) across the visible range (300–700 nm) (Fig. 2B). The PP morph exhibited low reflectance at shorter wavelengths with a pronounced increase beyond ∼600 nm, consistent with anthocyanin-dominated pigmentation. In contrast, YW and YY morphs showed elevated reflectance between ∼500–600 nm and high reflectance at longer wavelengths, consistent with reduced anthocyanin and increased carotenoid contributions. The RW morph displayed intermediate spectral characteristics, with elevated reflectance in the green–yellow region. These distinct and non-overlapping spectral profiles support classification of morphs as discrete phenotypic categories rather than continuous variation. The RWh morph was rare and restricted to sympatric zones between PP and RW populations. Although spectral measurements were not obtained due to limited sampling, representative inflorescences indicate a mosaic phenotype combining red floral tubes with mixed white–pink lobes (Fig. 2C–G), distinguishing it from both parental morphs.

To ensure that downstream expression analyses were not confounded by assembly or annotation bias, we evaluated transcriptome completeness and structural consistency across morphs and tissues. BUSCO analyses indicated uniformly high completeness (97–99%) with comparable duplication and fragmentation levels (Supplementary Fig. S1A). Expression-weighted contiguity (ExN50) increased with transcript abundance and reached ∼2.5 kb at Ex90 (Supplementary Fig. S1B), indicating high assembly quality. Orthogroup analysis revealed extensive transcript sharing among morphs, indicating a largely conserved gene repertoire (Supplementary Fig. S1C). Mean–variance relationships followed the expected negative binomial dispersion (Supplementary Fig. S1D), and transcript length and isoform distributions were comparable across datasets (Supplementary Fig. S1E–K). RNA libraries were of high quality (mapping rates ∼95–97%, Q30 >94%), and *Trinity* assemblies showed consistent contiguity and gene representation before and after filtering (Tables S2–S4). Together, these results confirm that transcriptomic datasets are comparable across morphs and suitable for downstream comparative analyses.

### Phylogenetic placement and genome-wide expression inheritance of the RWh morph

To evaluate whether the mosaic phenotype of the RWh morph reflects intermediate evolutionary ancestry, we reconstructed a morph-level phylogeny using genome-wide single-copy orthologs inferred with OrthoFinder. The resulting topology was well resolved and did not support a simple hybrid origin of RWh between the sympatric parental morphs (PP and RW) (Fig. 2H). Instead, RWh was nested within the PP lineage as its closest relative, with RW forming a more distant sister clade. This placement indicates that the mosaic phenotype of RWh does not correspond to phylogenetic intermediary between PP and RW. Consistent with this phylogenetic pattern, orthogroup analysis revealed that the vast majority of RWh genes were assigned to shared orthogroups (94%), with a low proportion of morph-specific genes (3.7%) relative to parental morphs (Supplementary Table S5). These results indicate that RWh does not represent a genomically distinct lineage but instead shares a largely conserved gene repertoire with parental morphs.

We next assessed whether gene expression in RWh follows additive expectations along the PP–RW axis using a hybrid index framework. Genome-wide classification revealed that intermediate expression was rare, representing only 6.89% of genes after scaling (Table 1). Parent-like expression patterns were also uncommon (PP-like: 4.93%; RW-like: 2.16%). In contrast, the majority of genes (86.02%) exhibited transgressive expression outside the parental range. This transgressive pattern was asymmetric, with most genes showing expression below PP levels (74.33%), whereas a smaller fraction exceeded RW expression (11.69%).

**Table 1.** Genome-wide inheritance patterns based on expression hybrid index.

| Category | Gene count |  |  |  |
| --- | --- | --- | --- | --- |
|  | Raw (n) | Raw (%) | Scaled (n) | Scaled (%) |
| Intermediate | 4,446 | 8.93 | 3,463 | 6.89 |
| PP-like | 3,530 | 7.09 | 2,476 | 4.93 |
| RW-like | 4,172 | 8.38 | 1,085 | 2.16 |
| Transgressive below PP | 24,643 | 49.50 | 37,343 | 74.33 |
| Transgressive above RW | 12,995 | 26.10 | 5,871 | 11.69 |
| Total transgressive | 37,638 | 75.60 | 43,214 | 86.02 |

Together, the phylogenetic placement and genome-wide expression analyses indicate that RWh does not represent a simple additive hybrid state. Instead, the mosaic phenotype is associated with extensive transgressive regulatory variation and phylogenetic affinity to the PP lineage. These results support a model in which RWh reflects a derived regulatory state characterized by widespread non-additive expression rather than intermediate inheritance.

### Gene expression divergence reveals canalized regulatory states and transgressive expansion

Genome-wide gene expression profiles were strongly structured by floral morph rather than geographic origin. Principal component analysis of normalized expression data separated individuals into discrete clusters corresponding to the five morphs, with PC1 and PC2 explaining 21% and 11% of total variance, respectively (Fig. 3A). Parental morphs formed compact, non-overlapping clusters, indicating low within-morph transcriptional variance and consistent regulatory states across populations. In contrast, the RWh morph occupied a displaced and expanded region of expression space relative to the parental axis, consistent with increased regulatory dispersion rather than simple additive inheritance.

**Figure 3.**
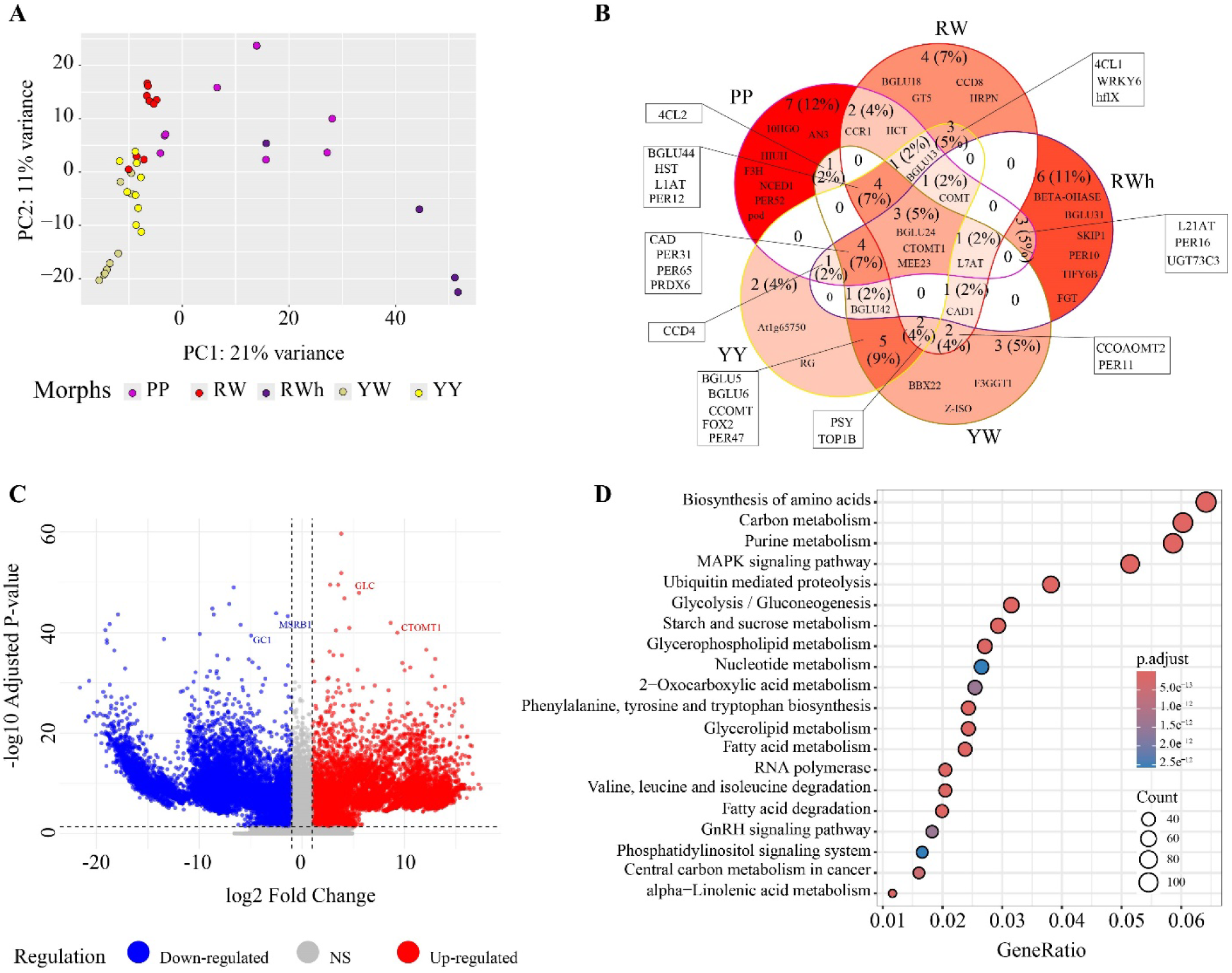
Gene expression divergence and functional differentiation among floral morphs. (**A**) Principal component analysis (PCA) of normalized gene expression profiles showing clustering by floral morph (PP, RW, RWh, YW, YY). Points represent individuals colored by morph. PC1 and PC2 explain 21% and 11% of total variance, respectively, revealing strong morph-associated transcriptional structure. (**B**) Venn diagram showing the number and overlap of morph-associated differentially expressed genes (DEGs) among morphs. Numbers indicate gene counts, with percentages relative to the total number of DEGs. Representative genes from key metabolic and regulatory pathways are indicated. (**C**) Volcano plot of differential gene expression based on a generalized linear model (see Methods). Each point represents a gene plotted by log₂ fold change and −log₁₀ adjusted p-value. Positive and negative log₂ fold change values indicate relatively higher expression in one or the other morph depending on the model contrast. Significantly up-regulated genes are shown in red, down-regulated genes in blue, and non-significant genes in gray. Pairwise comparisons are shown in Supplementary Fig. S2. (**D**) KEGG pathway enrichment of DEGs. Dot size represents gene counts and color indicates adjusted p-values. Enriched pathways include amino acid biosynthesis, carbon metabolism, signal transduction, and secondary metabolism associated with pigmentation and stress responses.

Differential expression analyses revealed extensive morph-associated transcriptional divergence (Fig. 3B–C; Supplementary Fig. S2). Across pairwise contrasts, thousands of genes were significantly differentially expressed (FDR-adjusted P < 0.05), with both shared and morph-specific components, with directionality dependent on the comparison order (i.e., relative up- or down-regulation between morphs; Supplementary Fig. S2). Although a small core set of DEGs was shared among morphs, substantial fractions were morph-specific, indicating that floral morphs correspond to discrete transcriptional programs rather than continuous variation.

The PP morph showed the largest fraction of uniquely expressed genes (∼12%), with consistent up-regulation of flavonoid and anthocyanin biosynthesis genes across multiple pairwise comparisons (e.g., PP vs RW, PP vs RWh, PP vs YW; Supplementary Fig. S2). Key structural genes, including *F3H, ANS, CCR1, HCT, 4CL2*, and *L1AT*, were repeatedly more highly expressed in PP relative to all other morphs (Fig. 3C; Supplementary Fig. S2; Supplementary Table S6), consistent with elevated anthocyanin flux underlying pink pigmentation. The PP morph also showed enrichment of regulatory and stress-associated genes such as *WRKY6, HST, PER12*, and *PRDX6*, suggesting coordinated regulation of pigment biosynthesis and redox-related processes. The RW morph exhibited a smaller but distinct set of DEGs (∼7%), including *BGLU18, CCD8, GT5,* and *HRPN*, which are associated with carotenoid cleavage, glycosylation, and secondary metabolite modification (Fig. 3B; Supplementary Table S6). Several phenylpropanoid-associated genes (e.g., *COMT, CTOMT1, BGLU24*) were shared between PP and RW, indicating partial overlap in regulatory architecture, as also reflected in their pairwise contrasts (Supplementary Fig. S2). Yellow-based morphs (YW and YY) displayed strongly divergent transcriptional profiles relative to PP and RW, with minimal overlap in DEGs across pairwise comparisons (Supplementary Fig. S2). The YY morph uniquely expressed genes involved in carotenoid metabolism and plastid function, including *BGLU5, BGLU6, CCOMT, FOX2*, and *PER47*, whereas the YW morph was characterized by genes such as *BBX22, Z-ISO, F3GGT1*, and *CCOAOMT2* (Fig. 3B; Supplementary Table S6). These patterns are consistent with pigmentation primarily driven by carotenoid accumulation and reduced anthocyanin contribution.

The RWh morph exhibited a mixed but non-additive expression profile. While subsets of genes displayed a distinct set of uniquely expressed genes (∼11%), including *BETA-OHASE*, *BGLU31, FGT, SKIP1, PER10*, and *TIFY6B*, a substantial fraction displayed transgressive patterns relative to both parental morphs (Fig. 3B; Supplementary Table S6). These genes are associated with hormone signaling, transcriptional regulation, and stress responses, indicating activation of regulatory pathways not predominant in either parent. Notably, several core anthocyanin biosynthesis genes (e.g., *ANS, F3H*) showed intermediate expression, whereas others (e.g., *COMT, CTOMT1*) were reduced relative to both parents. In contrast, genes associated with stress response and signaling, including *MSRB1*, *GC1*, and *NCED1*, were frequently up-regulated beyond parental levels, indicating activation of regulatory pathways not predominant in either parental morph.

Hierarchical clustering of pigment- and metabolism-associated genes further highlighted differences in regulatory stability among morphs (Supplementary Fig. S3). Parental morphs displayed highly coordinated expression patterns. Specifically, PP and RW consistently showed upregulation of key phenylpropanoid and flavonoid pathway genes (e.g., *HCT, BGLU13, L1AT, HST*), whereas YW and YY exhibited coordinated downregulation of these genes, accompanied by increased expression of carotenoid-associated genes such as *Z-ISO* and *F3GGT1*. In contrast, the RWh morph showed heterogeneous expression across these clusters, combining intermediate (*ANS, F3H*), parent-biased, and transgressive patterns (e.g., *COMT, PER12, BGLU24*). These results indicate that while parental morphs maintain coordinated regulatory programs, the RWh morph reflects partial disruption of these co-regulated gene networks.

### Functional enrichment reflects metabolic bases of morph differentiation

Gene Ontology (GO) enrichment analyses revealed functional categories consistent with the metabolic and regulatory basis of floral pigmentation (Supplementary Fig. S4). Across morphs, enriched biological processes included phenylpropanoid and secondary metabolite biosynthesis, oxidation–reduction processes, and responses to stimuli, reflecting the coordinated metabolic and regulatory demands of pigment production. Enrichment of plastid- and membrane-associated components further highlights the cellular context of carotenoid biosynthesis and metabolite transport.

Morph-specific enrichment patterns were consistent with distinct pigmentation strategies. The PP morph showed strong enrichment of flavonoid and phenylpropanoid pathways together with redox-related processes (Supplementary Fig. S4C), consistent with high anthocyanin flux and associated oxidative regulation. The RW morph was enriched for regulatory and signaling categories (Supplementary Fig. S4B), suggesting modulation of pigment pathways rather than full pathway activation. In contrast, yellow-based morphs (YW and YY) were enriched for plastid-associated metabolic processes (Supplementary Fig. S4E, F), consistent with carotenoid biosynthesis and storage within plastids. The RWh morph exhibited broader and less specialized enrichment across metabolic, regulatory, and stress-response categories (Supplementary Fig. S4D), indicating a more heterogeneous transcriptional state. This contrasts with the more functionally coherent enrichment profiles observed in parental morphs and is consistent with partial disruption of coordinated regulatory programs.

The KEGG pathway analyses further supported these patterns by highlighting differences in central metabolic allocation (Fig. 3D; Supplementary Fig. S5). Core pathways such as carbon metabolism, amino acid biosynthesis, and MAPK signaling were enriched across morphs, reflecting the metabolic cost and regulatory control of pigment production. However, pathway representation differed among morphs: PP and RW showed stronger enrichment for secondary metabolism and signaling pathways, whereas YW and YY were enriched for lipid metabolism and plastid-associated processes. Notably, the RWh morph showed the strongest enrichment of central metabolic pathways, including glycolysis/gluconeogenesis and fatty acid metabolism, suggesting reorganization of primary metabolic activity. Together, these results indicate that floral color polymorphism is embedded within broader metabolic and regulatory networks rather than driven by isolated pigment pathways.

### Modular regulatory architecture underlies floral morph differentiation

Weighted gene co-expression network analysis revealed a modular regulatory architecture underlying floral morph differentiation (Fig. 4A; Supplementary Fig. S6). Genes clustered into discrete co-expression modules of varying sizes, each representing coordinated transcriptional programs. The network construction achieved approximate scale-free topology at the selected soft-thresholding power, with a clear trade-off between scale-free model fit and mean connectivity (Supplementary Fig. S6A, B). Hierarchical clustering of genes based on topological overlap delineated well-defined modules of varying sizes, each summarized by a module eigengene (Supplementary Fig. S6C). These modules provide a systems-level framework for evaluating how gene expression is organized across morphs.

**Figure 4.**
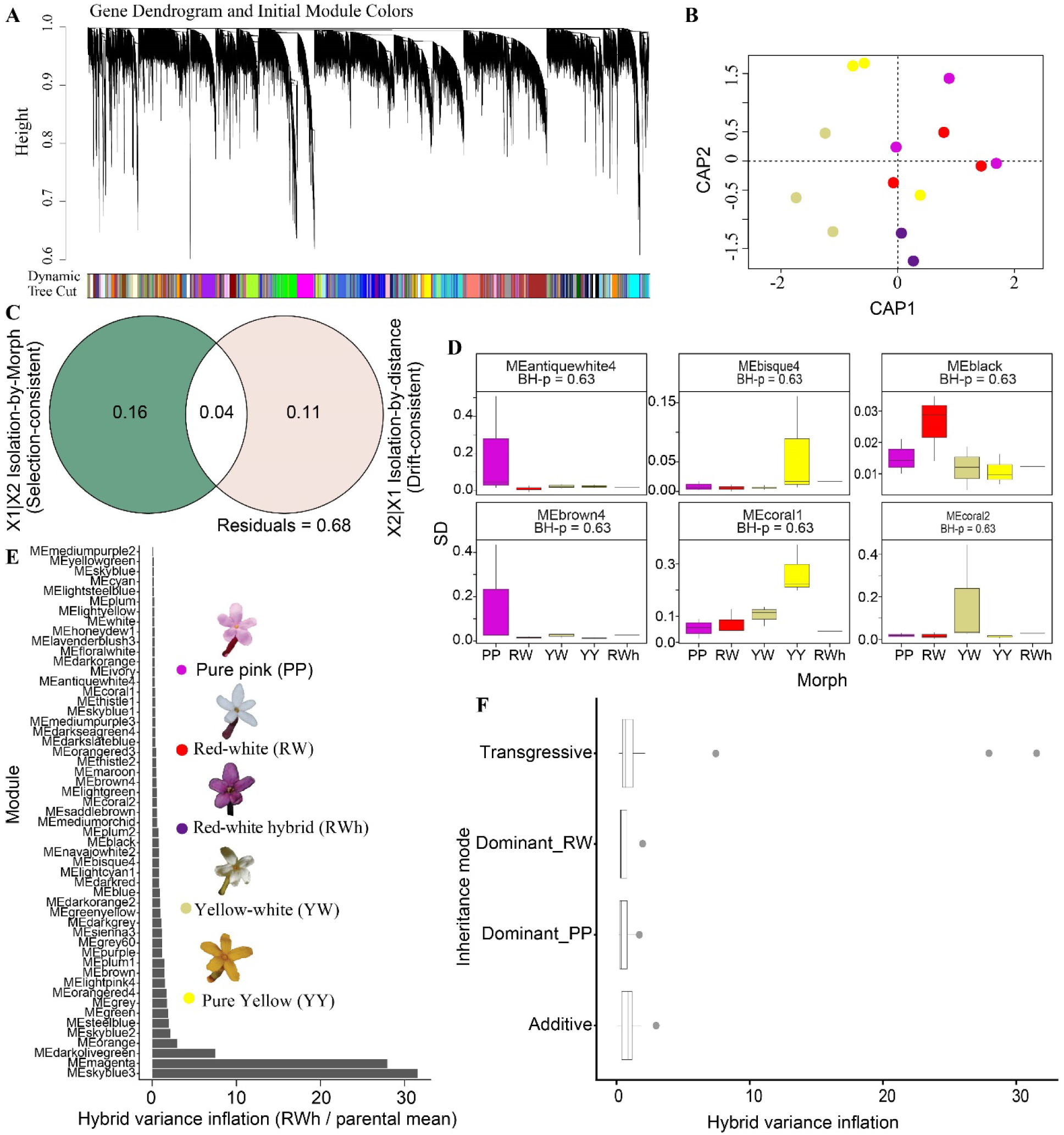
Canalization and modular destabilization of gene regulatory networks across floral morphs. (**A**) Hierarchical clustering of genes based on expression similarity, with co-expression modules identified using dynamic tree cutting. Colored bars indicate module assignments. (**B**) Canonical analysis of principal coordinates (CAP) based on population-mean module eigengenes. Points represent morph means (PP, RW, RWh, YW, YY), showing morph-associated structure in regulatory profiles. (**C**) Variance partitioning of module eigengene variation into components associated with morph identity, geographic structure, their shared contribution, and residual variance. (**D**) Within-morph expression variance (standard deviation, SD) across co-expression modules (n = 53). Boxplots show reduced variance in parental morphs and elevated variance in subsets of modules in RWh. Adjusted P-values (Benjamini–Hochberg) are indicated. (**E**) Hybrid variance inflation across modules, calculated as the ratio of variance in RWh relative to the parental mean. Modules are ordered by increasing inflation, showing that most remain stable while a subset exhibit pronounced increases. (**F**) Hybrid variance inflation across inheritance modes (additive, dominant PP, dominant RW, transgressive). Points represent individual modules, with higher variance inflation primarily associated with transgressive inheritance.

Module–trait correlation analyses identified strong and morph-specific regulatory associations (Supplementary Fig. S6D). Several modules were positively correlated with individual parental morphs, indicating that each morph is characterized by a distinct combination of regulatory programs rather than uniform shifts in gene expression. In contrast, modules associated with the RWh morph did not simply mirror parental patterns, suggesting that its regulatory organization is not intermediate at the network level. Hierarchical clustering of module eigengenes further revealed higher-order structure among modules, grouping regulatory programs with shared expression dynamics (Supplementary Fig. S6E). Sample clustering based on module eigengenes confirmed clear separation among morphs without evidence of outliers (Supplementary Fig. S6F), supporting the robustness of the inferred network structure. Functional enrichment of morph-associated modules indicated that regulatory architecture reflects biological specialization. Modules associated with PP and RW were enriched for phenylpropanoid and flavonoid biosynthesis pathways, whereas modules associated with YW and YY were enriched for carotenoid metabolism and plastid-associated processes (Supplementary Fig. S7). Modules associated with the RWh morph showed enrichment for regulatory and stress-response functions, consistent with altered transcriptional states relative to parental morphs. The absence of intermediate module-level patterns in RWh further indicates that regulatory variation in this morph reflects reorganization of network structure rather than additive inheritance.

### Canalization and selective hybrid destabilization shape regulatory divergence

To disentangle morph-associated regulatory divergence from neutral spatial structure, we partitioned variation in population-mean module eigengenes using PERMANOVA, distance-based redundancy analysis (dbRDA), and variance partitioning (Table 2). Canonical analysis of principal coordinates based on population-mean module eigengenes further separated morphs along major axes of regulatory variation, with the RWh morph positioned between PP and RW but shifted away from the direct parental axis, occupying a distinct region of expression space rather than a simple mid-parent position (Fig. 4B). Morph identity explained a substantial fraction of transcriptomic variation (R² = 0.45, P = 0.001), indicating strong phenotype-associated structuring of regulatory profiles. After accounting for spatial structure using geographic eigenvectors, morph identity retained a marginal but persistent effect (dbRDA: F = 1.43, P = 0.08), suggesting that morph-associated divergence is not fully explained by geographic distance alone (Fig. 4B; Table 2). Variance partitioning further revealed a larger unique contribution of morph identity (adjusted R² = 0.16) relative to geography (adjusted R² = 0.12), with a smaller shared component (adjusted R² = 0.05). Together, these results indicate that transcriptomic divergence aligns more strongly with morph identity than with spatial structure (Fig. 4C).

**Table 2.**
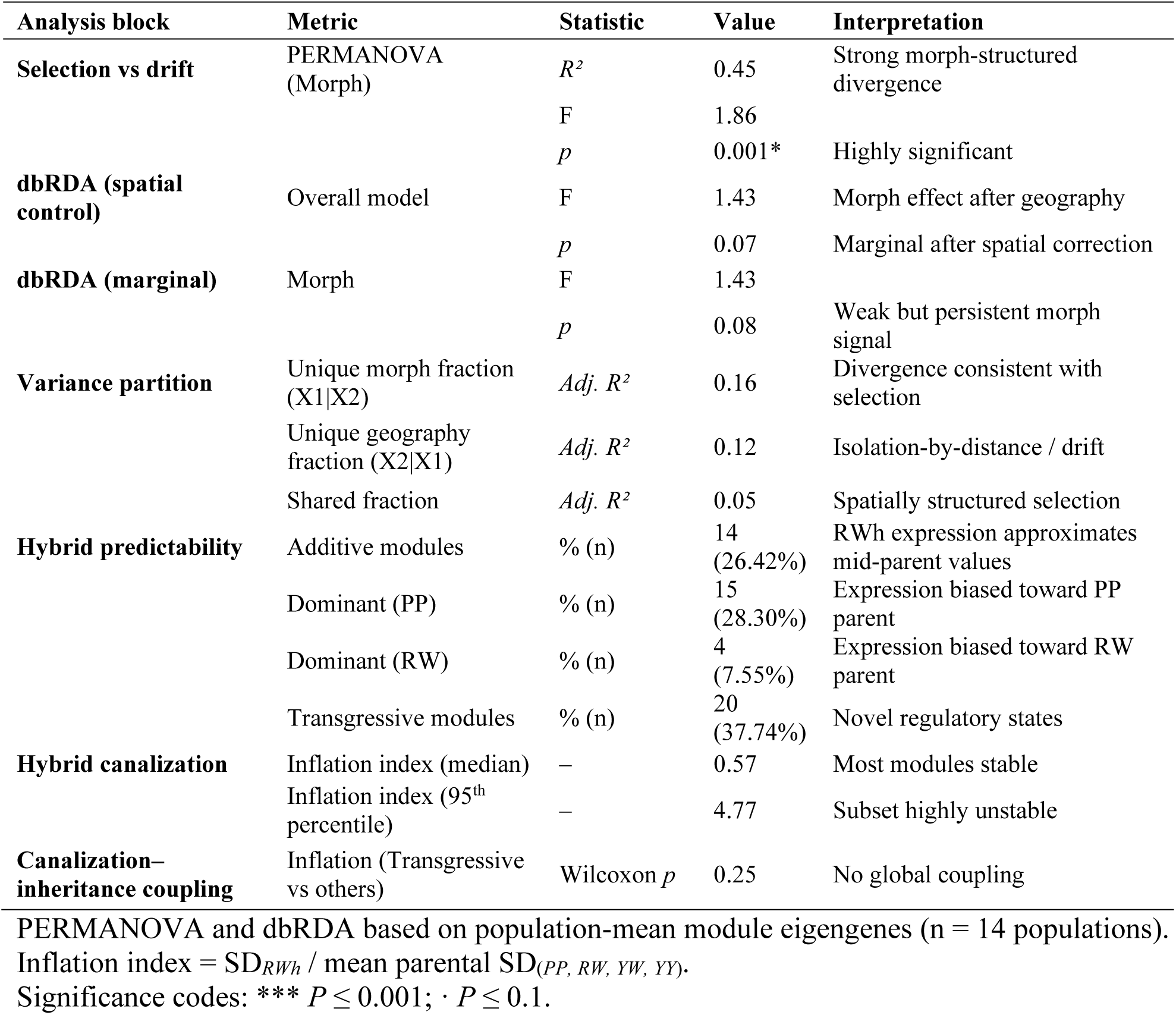
Eco-evolutionary architecture of transcriptomic divergence, canalization, and RWh expression predictability.

Within-population variance of module eigengenes provided a quantitative measure of transcriptional constraint across morphs (Fig. 4D; Supplementary Fig. S8A). Parental morphs (PP, RW, YW, YY) consistently exhibited low variance across modules, indicating stable and tightly regulated expression states. In contrast, the RWh morph showed a similar median variance but a pronounced tail of high-variance modules, indicating elevated variability in a subset of regulatory programs.

Hybrid variance inflation analysis further quantified this pattern. The median inflation index across modules was low (0.57), indicating that most regulatory modules remained stable in RWh relative to parental morphs (Table 2). However, a small subset of modules exhibited extreme inflation, with values exceeding 25 in modules such as MEskyblue3 and MEmagenta (Table 3). These modules (arbitrarily labeled by WGCNA) represent distinct gene regulatory subnetwork, suggesting that regulatory instability is localized to specific functional pathways rather than diffusely distributed across the transcriptome (Fig. 4E; Table 2). Module-level inheritance analysis revealed structured but heterogeneous regulatory outcomes in RWh (Fig. 4F; Supplementary Fig. S8B; Table 3). Across all modules (n = 53), 26.4% (14 modules) exhibited additive inheritance, while 35.8% (28.3% dominant-PP and 7.5% dominant-RW) showed dominant patterns biased toward one parental morph. In contrast, 37.7% of modules (20 modules) displayed transgressive inheritance, with expression extending beyond the parental range, consistent with the emergence of novel regulatory states (Fig. 4F; Table 2).

**Table 3.** Top five transcriptional modules showing the strongest hybrid variance inflation and distinct inheritance modes.

| Module | Inheritance mode | SD<sub>PP</sub> | SD<sub>RW</sub> | SD<sub>RWh</sub> | SD<sub>YW</sub> | SD<sub>YY</sub> | Parental mean SD | Inflation index |
| --- | --- | --- | --- | --- | --- | --- | --- | --- |
| MEskyblue3 | Transgressive | 0.0273 | 0.0134 | 0.4730 | 0.0106 | 0.0087 | 0.0150 | 31.52 |
| MEmagenta | Transgressive | 0.0351 | 0.0087 | 0.4420 | 0.0047 | 0.0147 | 0.0158 | 27.90 |
| MEdarkolivegreen | Transgressive | 0.0411 | 0.0309 | 0.2310 | 0.0256 | 0.0259 | 0.0309 | 7.47 |
| MEorange | Dominant_PP | 0.0365 | 0.0233 | 0.0680 | 0.0108 | 0.0206 | 0.0228 | 2.98 |
| MEskyblue2 | Additive | 0.0545 | 0.0175 | 0.0820 | 0.0458 | 0.0341 | 0.0380 | 2.15 |

Hybrid variance inflation indices were calculated for the transgressive morph (*RWh*) relative to the mean within-population variance of the two closest parental morphs (*PP* and *RW*). Modules were classified as additive, dominant, or transgressive based on hybrid module eigengene means relative to parental morphs. Standard Deviation (SD) values represent within-population standard deviations of module eigengenes.
Inflation index = $SD_{RWh} / \text{mean parental } SD_{(PP, RW, YW, YY)}$ ; Parental morphs (*PP* and *RW*) were identified as the closest parental types to *RWh* based on module-level expression distance.

Modules exhibiting transgressive inheritance tended to show higher variance inflation than additive or dominant modules, although this relationship was not statistically significant (Wilcoxon P = 0.25; Table 2). The most extreme cases of hybrid regulatory disruption were confined to a few modules; for example, MEskyblue3 showed a hybrid variance inflation index of 31.52 together with transgressive inheritance. Together, these results indicate that most gene regulatory modules remain stable across morphs, whereas a subset is selectively destabilized in the RWh morph. This pattern supports a model in which transcriptional systems are largely robust but permit localized breakdown of regulatory constraint, generating transgressive expression and potential phenotypic novelty.

## Discussion

### Morph identity defines stable transcriptional states across geographic space

The central goal of this study was to determine whether flower-color polymorphism in *Stellera chamaejasme* reflects transient phenotypic variation or stable regulatory states associated with morph identity. Conceptually, transient phenotypic variation would be expected to produce weak and inconsistent transcriptional differentiation among morphs, with expression patterns primarily structured within populations or across environmental gradients. In contrast, persistent morph-associated regulatory states would be expected to show stronger and more consistent transcriptional divergence among morphs than within morphs or populations, remain stable across geographic space, and persist after accounting for spatial structure. Across populations spanning broad elevational gradients (2,676–4,210 m a.s.l.) in Southwest China, transcriptomic variation was consistently structured by morph rather than geographic distance. Partitioning analyses further showed that morph identity explained a larger and distinct component of expression variation than geography, and this signal remained after explicit spatial correction. These results indicate that floral morphs correspond to reproducible phenotype-associated transcriptional programs and reveal an isolation-by-morph pattern at the regulatory level.

Disentangling phenotype-associated divergence from spatial structure is particularly challenging in topographically complex regions such as the Qinghai–Tibetan Plateau, where environmental and geographic gradients are strongly correlated (Meirmans, 2012; Wang & Bradburd, 2014). By explicitly modeling spatial effects, our analyses provide a conservative test of morph-associated divergence and reduce the likelihood of spurious inference. The persistence of morph-associated transcriptional structure after spatial correction indicates that regulatory differentiation is unlikely to arise solely from isolation by distance or demographic history.

Although transcriptomic data alone cannot directly demonstrate the mode of selection, the observed patterns are consistent with morph-specific regulatory configurations potentially maintained by phenotype-associated or selective processes. Comparable phenotype-linked regulatory differentiation has been reported in several floral polymorphism systems, including monkeyflowers *Mimulus* L. (Schemske & Bradshaw, 1999)/*Mimulus sect. Erythranthe* (Spach) Greene (Wenzell *et al*., 2025), *Penstemon* Schmidel (Wessinger *et al*., 2014), and *Phlox* L. (Hopkins & Rausher, 2012), as well as in eco-genomic studies showing that phenotypic predictors can explain expression or genomic variation beyond spatial structure (Wang & Bradburd, 2014; Forester *et al*., 2016; Capblancq *et al*., 2020). The shared morph–geography component observed here further suggests that environmental gradients may also influence morph distributions through variation in abiotic conditions, pollinator communities, or resource availability. Such coupling is expected when selection varies along environmental gradients but favors discrete alternative phenotypes rather than continuous variation (Schemske & Bradshaw, 1999; Hopkins & Rausher, 2012). Importantly, the substantial morph-specific component independent of geography suggests that regulatory divergence is associated with morph identity itself rather than arising solely as a by-product of environmental variation. Collectively, these findings indicate that flower-color morphs in *S. chamaejasme* represent stable regions of transcriptional state space that recur across heterogeneous landscapes.

### Canalization of regulatory networks underlies morph stability

Canalization is a key feature of stabilizing selection, where phenotypic and gene expression variation is reduced around an optimal regulatory state (Waddington, 1942; Gibson & Wagner, 2000; Masel & Siegal, 2009). Consistent with this expectation, parental morphs exhibited low within-population and within-morph variance in co-expression module eigengenes, even among geographically separated populations spanning approximately 23–410 km. Notably, this pattern is observed at the level of regulatory modules rather than individual genes, suggesting that morph differentiation is embedded within network-level expression architectures. Such module-level stability reflects regulatory systems that buffer expression against environmental and genetic perturbation.

Module-level canalization reflects regulatory systems that buffer expression against environmental and genetic perturbation (Rifkin *et al*., 2005; Hodgins-Davis & Townsend, 2009; McManus *et al*., 2010). The persistence of these low-variance expression regimes across geographically separated populations suggests that each floral morph occupies a distinct and stable region of transcriptional space consistent with maintenance by selective or ecological filtering processes. Independent population genomic evidence showing strong regional structure and limited gene flow further indicates that these regulatory configurations persist despite demographic differentiation (Rana *et al*., 2024).

This pattern was particularly evident in pigmentation-associated regulatory modules. Genes involved in phenylpropanoid, flavonoid, and carotenoid metabolism were consistently organized into morph-specific co-expression modules exhibiting low within-morph variance. Because floral coloration depends on coordinated flux through multi-step biosynthetic pathways, robustness is expected to emerge at the pathway and network levels rather than at individual loci (Grotewold, 2006; Davies *et al*., 2012; Albert *et al*., 2014). The stability of these pigment-associated regulatory modules across populations therefore provides a direct mechanistic link between transcriptional canalization and the persistence of discrete flower-color phenotypes, reinforcing the view that morph differentiation in *S. chamaejasme* is deeply embedded in regulatory architecture rather than driven by isolated pigment genes (Rausher, 2008; Yuan *et al*., 2013; Monniaux, 2023).

### Transgressive morphs reveal limits of regulatory stability

The naturally occurring RWh morph provides insight into the limits of these regulatory states. Although field observations suggested a mosaic phenotype in zones of sympatry between PP and RW populations, ortholog-based phylogenetic and expression analyses do not support a simple additive hybrid interpretation. Instead, genome-wide hybrid index analyses revealed pervasive non-additive regulation, with approximately 86% of genes exhibiting expression outside the parental range. These results indicate that RWh represents a transgressive regulatory morphotype rather than an intermediate expression state.

When parental morphs correspond to stabilized regulatory configurations, recombination or regulatory reorganization may disrupt co-adapted interactions among network components, exposing epistatic interactions and relaxing stabilizing constraints (Muller, 1942; Dobzhansky, 1982; Rieseberg *et al*., 1999). Consistent with this pattern, the RWh morph exhibited elevated expression variance in a subset of co-expression modules. Importantly, this destabilization was modular rather than genome-wide. Most modules retained parental-like stability and predictable inheritance, whereas a limited subset showed pronounced variance inflation and transgressive expression. This selective destabilization aligns with theoretical and empirical evidence that regulatory robustness is unevenly distributed across gene networks, where core modules remain buffered while others are more sensitive to perturbation (Gibson & Wagner, 2000; Wagner, 2008; Paaby & Rockman, 2014), and is further supported by pathway-specific hybrid misexpression across diverse taxa (McManus *et al*., 2010; Mack & Nachman, 2017; Runemark *et al*., 2025).

Module-level inheritance patterns further indicate that regulatory novelty arises through localized disruption of network organization. While many modules exhibited additive or dominant expressions, a substantial proportion showed transgressive regulation beyond the parental range. Such expression patterns are commonly interpreted as signatures of epistatic interactions, compensatory regulatory evolution, or breakdown of co-adapted regulatory complexes (Rieseberg *et al*., 1999; Stelkens & Seehausen, 2009; Camper *et al*., 2025; Runemark *et al*., 2025). In *S. chamaejasme*, the coexistence of stable regulatory modules alongside a subset of transgressive modules suggests that gene regulatory networks can remain globally robust while permitting localized release of expression variation. Notably, several of the modules exhibiting transgressive regulation were enriched for pigment biosynthesis and secondary metabolism, directly linking reduced regulatory constraint to floral color variation.

### Regulatory integration links flower pigmentation to broader physiological organization

Our results further indicate that flower-color morphs in *S. chamaejasme* are not defined solely by pigment biosynthesis but correspond to coordinated transcriptional states involving multiple physiological processes. Functional enrichment analyses revealed that morph-associated expression divergence extends beyond anthocyanin and flavonoid pathways to include carbon metabolism, redox processes, hormone signaling, and developmental regulation. These findings indicate that flower-color polymorphisms reflect integrated physiological states rather than isolated biochemical traits.

This integrative organization is increasingly recognized as a defining feature of floral trait evolution. Pigment pathways are tightly linked to primary metabolism, stress responses, and developmental regulation (Smith & Rausher, 2011; Martínez-Harms *et al*., 2022; Yeo & Moyroud, 2025). Anthocyanins, for example, contribute not only to pollinator attraction but also to UV protection, thermal regulation, and oxidative stress mitigation, particularly in high-elevation environments (Neill & Gould, 2003; Davies *et al*., 2012; Landi *et al*., 2015). Because these functions share metabolic substrates and regulatory control, selection acting on ecological performance may indirectly contribute to stabilizing pigment-associated regulatory programs (Paaby & Rockman, 2014; Wessinger & Rausher, 2015). In this context, module-level canalization may contribute to maintaining functional integration by ensuring that pigmentation remains coordinated with broader physiological processes (Gibson & Wagner, 2000; Masel & Siegal, 2009; Martínez-Harms *et al*., 2022), whereas disruption of these modules can generate both expression variability and phenotypic novelty. The transgressive RWh morph illustrates this pattern, showing altered regulation in modules enriched for metabolic and regulatory functions, suggesting partial decoupling of coordinated gene networks (Landry *et al*., 2007; McManus *et al*., 2010; Mack & Nachman, 2017). In *S. chamaejasme*, this pattern implies that regulatory integration both stabilizes discrete floral phenotypes and constrains the range of viable transgressive expression states.

### Evolutionary implications and future directions

Together, our results indicate that flower-color morphs in *S. chamaejasme* correspond to discrete and repeatable transcriptional states characterized by low within-morph variance and strong phenotype-associated structure across populations. These patterns are consistent with morph-specific regulatory configurations potentially maintained by stabilizing or environmentally mediated processes rather than neutral demographic structure alone. The presence of extensive transgressive expression and localized variance inflation in the RWh morph further suggests that recombination or regulatory reorganization may expose incompatibility among co-adapted network components, generating phenotypic novelty through localized disruption of regulatory buffering (Seehausen *et al*., 2014).

Several limitations expands from considering sampling strategy to the fitness assessment. Transcriptomic analyses were restricted to a specific tissue and developmental stage, and the strength of canalization may vary across ontogeny or environmental conditions. Sampling of the transgressive morph was limited due to its rarity, which may reduce power to detect subtle effects. In addition, while the observed patterns are consistent with regulatory constraint, direct tests of selection and fitness consequences will be required to establish the evolutionary mechanisms underlying morph persistence.

Future work linking regulatory architecture to ecological performance will be essential for evaluating the evolutionary consequences of these transcriptional states. Experimental approaches such as pollinator preference assays, reciprocal transplant experiments, and functional validation of regulatory loci could clarify the fitness consequences of alternative morph-specific regulatory programs. Integrating transcriptomic architecture with population genomic and ecological data will be necessary to understand how regulatory robustness shapes the persistence, breakdown, and evolutionary potential of polymorphic traits in natural populations.

## Supporting information

Supplementary Informations

## Acknowledgements

We thank Zhen-Yu Lv for assistance with field sampling, Dr. Zhe Chen for taking the reflectance measurements, and Ms. Song Min-shu for laboratory support. This work was supported by the Key Projects of the Joint Fund of the National Natural Science Foundation of China (U23A20149), Yunnan Key R&D Program (202403AC00028), the Second Tibetan Plateau Scientific Expedition and Research (STEP) Program (2024QZKK0200), the Research Fund for International Young Scientists of the National Natural Science Foundation of China (32150410356), the National Natural Science Foundation of China (32322006), and the Young Academic and Technical Leader Raising Foundation of Yunnan Province (2019HB039). The first author was additionally supported by the CAS President’s International Fellowship Initiative (PIFI) postdoctoral fellowship (2021PB0034).

## Author contributions

S.K.R., T.D. and H.S. planned and designed the research. S.K.R., H.K.R. and C.J. performed the research and conducted field and laboratory work. S.K.R. and H.K.R. performed transcriptome assembly and analysed the data. S.K.R. conducted eco-evolutionary and network analyses. S.K.R. and H.K.R. prepared the figures. T.D. and H.S. supervised the project and acquired funding. S.K.R. wrote the manuscript, and all authors contributed to reviewing, editing, and approving the final version.

## Data availability

Raw RNA-seq reads generated in this study were deposited in the NCBI Sequence Read Archive (SRA) under BioProject accession PRJNA1451939. Processed data, including expression matrices, differential expression results, co-expression modules, and functional annotation outputs, are available in the GitHub repository: https://github.com/santoshkumarrana/stellera-regulatory-network-polymorphism and archived in Zenodo (10.5281/zenodo.19307586) (Rana, 2026). All data will be publicly accessible upon publication.

## Code availability

Custom scripts developed for this manuscript are publicly available on GitHub: https://github.com/santoshkumarrana/Canalized-regulatory-networks-stabilize-floral-polymorphism-and-enable-localized-transgression. A version of the repository has been archived at Zenodo (10.5281/zenodo.19307586) (Rana, 2026) to ensure long-term accessibility.

## Supporting Information (brief legends)

## Supplementary Figures

**Supplementary Fig. S1** Transcriptome assembly quality, expression characteristics, and transcript diversity.

**Supplementary Fig. S2** Pairwise differential gene expression among flower-color morphs.

**Supplementary Fig. S3** Gene expression heatmap for pigment- and metabolism-associated genes.

**Supplementary Fig. S4** Gene Ontology enrichment of differentially expressed genes across flower-color morphs.

**Supplementary Fig. S5** KEGG pathway enrichment of differentially expressed genes among flower-color morphs.

**Supplementary Fig. S6** Weighted gene co-expression network analysis (WGCNA) identifies modules associated with flower-color morphs.

**Supplementary Fig. S7** Pathway representation and functional network structure of pigment-associated genes.

**Supplementary Fig. S8** Within-morph expression variance and inheritance patterns of co-expression modules.

## Supplementary Tables

**Supplementary Table S1** Population sampling, geographic coordinates, and experimental design.

**Supplementary S2** Basic information of the population’s vouchers, transcriptome sequencing, and alignment rates of RNA libraries of four tissues of four different morphs of *Stellera chamaejasme*.

**Supplementary S3** Statistics of the Trinity assembly of transcriptomes for four sample tissues of four different morphs of *Stellera chamaejasme*.

**Supplementary Table S4** Statistics of the filtering of transcriptomes/pan- transcriptomes for four sample tissues of four different morphs of *Stellera chamaejasme*.

**Supplementary Table S5** OrthoFinder summary statistics for transcriptome assemblies across floral morphs of *Stellera chamaejasme*.

**Supplementary Table S6** Representative morph-associated DEGs and their functional roles in pigment biosynthesis and regulatory processes.

## References

Albert NW, Davies KM, Schwinn KE. 2014. Gene regulation networks generate diverse pigmentation patterns in plants. Plant Signaling & Behavior 9: e29526.

Benjamini Y, Hochberg Y. 1995. Controlling the false discovery rate: A practical and powerful approach to multiple testing. Journal of the Royal Statistical Society Series B: Statistical Methodology 57: 289–300.

Bolger AM, Lohse M, Usadel B. 2014. Trimmomatic: a flexible trimmer for Illumina sequence data. Bioinformatics 30: 2114–2120.

Borcard D, Gillet F, Legendre P. 2018. Numerical Ecology with R. Cham: Springer International Publishing.

Borcard D, Legendre P, Drapeau P. 1992. Partialling out the spatial component of ecological variation. Ecology 73: 1045–1055.

Boyle EA, Li YI, Pritchard JK. 2017. An expanded view of complex traits: From polygenic to omnigenic. Cell 169: 1177–1186.

Camacho C, Coulouris G, Avagyan V, Ma N, Papadopoulos J, Bealer K, Madden TL. 2009. BLAST+: Architecture and applications. BMC Bioinformatics 10: 421.

Camper BT, Kanes AS, Laughlin ZT, Manuel RT, Bewick SA. 2025. Transgressive hybrids as hopeful holobionts. Microbiome 13: 19.

Cantalapiedra CP, Hernández-Plaza A, Letunic I, Bork P, Huerta-Cepas J. 2021. eggNOG-mapper v2: Functional annotation, orthology assignments, and domain prediction at the metagenomic scale (K Tamura, Ed.). Molecular Biology and Evolution 38: 5825–5829.

Capblancq T, Fitzpatrick MC, Bay RA, Exposito-Alonso M, Keller SR. 2020. Genomic prediction of (mal)adaptation across current and future climatic landscapes. Annual Review of Ecology, Evolution, and Systematics 51: 245–269.

Chen H, Berg CS, Samuli M, Sotola VA, Sweigart AL, Yuan Y, Fishman L. 2025. The genetic architecture of floral trait divergence between hummingbird- and self-pollinated monkeyflower (*Mimulus*) species. New Phytologist 245: 2255–2267.

Dauphin B, Rellstab C, Wüest RO, Karger DN, Holderegger R, Gugerli F, Manel S. 2023. Re-thinking the environment in landscape genomics. Trends in Ecology & Evolution 38: 261–274.

Davies KM, Albert NW, Schwinn KE. 2012. From landing lights to mimicry: the molecular regulation of flower colouration and mechanisms for pigmentation patterning. Functional Plant Biology 39: 619–638.

Dobzhansky T. 1982. Genetics and the origin of species. New York: Columbia University Press.

Emms DM, Kelly S. 2019. OrthoFinder: Phylogenetic orthology inference for comparative genomics. Genome Biology 20: 238.

Félix M-A, Barkoulas M. 2015. Pervasive robustness in biological systems. Nature Reviews Genetics 16: 483–496.

Forester BR, Jones MR, Joost S, Landguth EL, Lasky JR. 2016. Detecting spatial genetic signatures of local adaptation in heterogeneous landscapes. Molecular Ecology 25: 104–120.

Forester BR, Lasky JR, Wagner HH, Urban DL. 2018. Comparing methods for detecting multilocus adaptation with multivariate genotype–environment associations. Molecular Ecology 27: 2215–2233.

Fu L, Niu B, Zhu Z, Wu S, Li W. 2012. CD-HIT: Accelerated for clustering the next-generation sequencing data. Bioinformatics 28: 3150–3152.

Gibson G, Wagner G. 2000. Canalization in evolutionary genetics: A stabilizing theory? BioEssays 22: 372–380.

Grabherr MG, Haas BJ, Yassour M, Levin JZ, Thompson DA, Amit I, Adiconis X, Fan L, Raychowdhury R, Zeng Q, et al. 2011. Full-length transcriptome assembly from RNA-Seq data without a reference genome. Nature Biotechnology 29: 644–652.

Gray SM, McKinnon JS. 2007. Linking color polymorphism maintenance and speciation. Trends in Ecology & Evolution 22: 71–79.

Grotewold E. 2006. The genetics and biochemistry of floral pigments. Annual Review of Plant Biology 57: 761–780.

Haas BJ, Papanicolaou A, Yassour M, Grabherr M, Blood PD, Bowden J, Couger MB, Eccles D, Li B, Lieber M, et al. 2013. De novo transcript sequence reconstruction from RNA-seq using the Trinity platform for reference generation and analysis. Nature Protocols 8: 1494–1512.

Hodgins-Davis A, Townsend JP. 2009. Evolving gene expression: from G to E to G×E. Trends in Ecology & Evolution 24: 649–658.

Hopkins R, Rausher MD. 2012. Pollinator-mediated selection on flower color allele drives reinforcement. Science 335: 1090–1092.

Horta-Lacueva QJ-B, Jónsson ZO, Thorholludottir DAV, Hallgrímsson B, Kapralova KH. 2023. Rapid and biased evolution of canalization during adaptive divergence revealed by dominance in gene expression variability during Arctic charr early development. Communications Biology 6: 897.

Kadelka C. 2026. Canalization as a stabilizing principle of gene regulatory networks: A discrete dynamical systems perspective. NPJ Systems Biology and Applications 12: 30.

Kadelka C, Murrugarra D. 2024. Canalization reduces the nonlinearity of regulation in biological networks. NPJ Systems Biology and Applications 10: 67.

Kanehisa M. 2000. KEGG: Kyoto encyclopedia of genes and genomes. Nucleic Acids Research 28: 27–30.

Kitano J, Kagawa K, Tsuchimatsu T, Yamaguchi R, Yamamichi M. 2025. The genomics of discrete polymorphisms maintained by disruptive selection. Trends in Ecology & Evolution 40: 1023–1034.

Landi M, Tattini M, Gould KS. 2015. Multiple functional roles of anthocyanins in plant-environment interactions. Environmental and Experimental Botany 119: 4–17.

Landry CR, Hartl DL, Ranz JM. 2007. Genome clashes in hybrids: insights from gene expression. Heredity 99: 483–493.

Langfelder P, Horvath S. 2008. WGCNA: An R package for weighted correlation network analysis. BMC Bioinformatics 9: 559.

Li J, Wang S, Miehe G, Opgenoorth L, Yang H, Wu D, Mao K. 2025. Ecological selection as drivers during early speciation: Insights from two allopatric cypress species in the Himalaya. Molecular Ecology 34: e70072.

Love MI, Huber W, Anders S. 2014. Moderated estimation of fold change and dispersion for RNA-seq data with DESeq2. Genome Biology 15: 550.

Mack KL, Nachman MW. 2017. Gene regulation and speciation. Trends in Genetics 33: 68–80.

Maia R, Gruson H, Endler JA, White TE. 2019. PAVO 2: New tools for the spectral and spatial analysis of colour in R (RB O’Hara, Ed.). Methods in Ecology and Evolution 10: 1097–1107.

Martínez-Harms J, Guerrero PC, Martínez-Harms MJ, Poblete N, González K, Stavenga DG, Vorobyev M. 2022. Mechanisms of flower coloring and eco-evolutionary implications of massive blooming events in the Atacama Desert. Frontiers in Ecology and Evolution 10: 957318.

Masel J, Siegal ML. 2009. Robustness: mechanisms and consequences. Trends in Genetics 25: 395–403.

McManus CJ, Coolon JD, Duff MO, Eipper-Mains J, Graveley BR, Wittkopp PJ. 2010. Regulatory divergence in *Drosophila* revealed by mRNA-seq. Genome Research 20: 816–825.

Meirmans PG. 2012. The trouble with isolation by distance. Molecular Ecology 21: 2839–2846.

Monniaux M. 2023. Unusual suspects in flower color evolution. Science 379: 534–535.

Muller HJ. 1942. Isolating mechanisms, evolution, and temperature.

Neill SO, Gould KS. 2003. Anthocyanins in leaves: Light attenuators or antioxidants? Functional Plant Biology 30: 865–873.

Oksanen J, Blanchet FG, Kindt R, Legendre P, Friendly M, McGlinn D, Minchin PR, O’Hara RB, Simpson GL, Solymos P, et al. 2019. vegan: Community ecology package.

Paaby AB, Rockman MV. 2014. Cryptic genetic variation: evolution’s hidden substrate. Nature Reviews Genetics 15: 247–258.

Patro R, Duggal G, Love MI, Irizarry RA, Kingsford C. 2017. Salmon provides fast and bias-aware quantification of transcript expression. Nature Methods 14: 417–419.

Payne JL, Wagner A. 2019. The causes of evolvability and their evolution. Nature Reviews Genetics 20: 24–38.

Peres-Neto PR, Legendre P, Dray S, Borcard D. 2006. Variation partitioning of species data matrices: Estimation and comparison of fractions. Ecology 87: 2614–2625.

Rana SK. 2026. Canalized gene regulatory networks stabilize floral polymorphism and enable modular transgressive expression in *Stellera chamaejasme*. v1.0.0, Zenodo, 29 Mar. 2026, 10.5281/zenodo.19307586

Rana SK, Rana HK, Landis JB, Kuang T, Chen J, Wang H, Deng T, Davis CC, Sun H. 2024. Pleistocene glaciation advances the cryptic speciation of *Stellera chamaejasme* L. in a major biodiversity hotspot. Journal of Integrative Plant Biology 66: 1192–1205.

Rausher MD. 2008. Evolutionary transitions in floral color. International Journal of Plant Sciences 169: 7–21.

Ravinet M, Faria R, Butlin RK, Galindo J, Bierne N, Rafajlović M, Noor MAF, Mehlig B, Westram AM. 2017. Interpreting the genomic landscape of speciation: a road map for finding barriers to gene flow. Journal of Evolutionary Biology 30: 1450–1477.

Rellstab C, Gugerli F, Eckert AJ, Hancock AM, Holderegger R. 2015. A practical guide to environmental association analysis in landscape genomics. Molecular Ecology 24: 4348–4370.

Rieseberg LH, Archer MA, Wayne RK. 1999. Transgressive segregation, adaptation and speciation. Heredity 83: 363–372.

Rifkin SA, Houle D, Kim J, White KP. 2005. A mutation accumulation assay reveals a broad capacity for rapid evolution of gene expression. Nature 438: 220–223.

Robinson MD, McCarthy DJ, Smyth GK. 2010. edgeR: A Bioconductor package for differential expression analysis of digital gene expression data. Bioinformatics 26: 139–140.

Romero IG, Ruvinsky I, Gilad Y. 2012. Comparative studies of gene expression and the evolution of gene regulation. Nature Reviews Genetics 13: 505–516.

Runemark A, Moore EC, Larson EL. 2025. Hybridization and gene expression: Beyond differentially expressed genes. Molecular Ecology 34: e17303.

Schemske DW, Bradshaw HD. 1999. Pollinator preference and the evolution of floral traits in monkeyflowers (*Mimulus*). Proceedings of the National Academy of Sciences 96: 11910–11915.

Seehausen O, Butlin RK, Keller I, Wagner CE, Boughman JW, Hohenlohe PA, Peichel CL, Saetre G-P, Bank C, Brännström Å, et al. 2014. Genomics and the origin of species. Nature Reviews Genetics 15: 176–192.

Simão FA, Waterhouse RM, Ioannidis P, Kriventseva EV, Zdobnov EM. 2015. BUSCO: assessing genome assembly and annotation completeness with single-copy orthologs. Bioinformatics 31: 3210–3212.

Smith SD. 2016. Pleiotropy and the evolution of floral integration. New Phytologist 209: 80–85.

Smith SD, Rausher MD. 2011. Gene loss and parallel evolution contribute to species difference in flower color. Molecular Biology and Evolution 28: 2799–2810.

Stelkens R, Seehausen O. 2009. Genetic distance between species predicts novel trait expression in their hybrids. Evolution 63: 884–897.

Sun S-F, Li Y-B, Leng J, Zhang D-H, Zhang W-L, Zhang B. 2024. Multiple insect pollination contributes to differential phenotypic selection on floral traits in *Stellera chamaejasme*. BMC Plant Biology 24: 1141.

Tigano A, Friesen VL. 2016. Genomics of local adaptation with gene flow. Molecular Ecology 25: 2144–2164.

Waddington CH. 1942. Canalization of development and the inheritance of acquired characters. Nature 150: 563–565.

Wagner A. 2008. Robustness and evolvability: A paradox resolved. Proceedings of the Royal Society B: Biological Sciences 275: 91–100.

Wagner A. 2011. The origins of evolutionary innovations: a theory of transformative change in living systems. Oxford: Oxford University Press.

Wang IJ, Bradburd GS. 2014. Isolation by environment. Molecular Ecology 23: 5649–5662.

Wenzell KE, Neequaye M, Paajanen P, Hill L, Brett P, Byers KJRP. 2025. Within-species floral evolution reveals convergence in adaptive walks during incipient pollinator shift. Nature Communications 16: 2721.

Wessinger CA, Hileman LC, Rausher MD. 2014. Identification of major quantitative trait loci underlying floral pollination syndrome divergence in *Penstemon*. Philosophical Transactions of the Royal Society B: Biological Sciences 369: 20130349.

Wessinger CA, Rausher MD. 2015. Ecological transition predictably associated with gene degeneration. Molecular Biology and Evolution 32: 347–354.

Wickham H, Averick M, Bryan J, Chang W, McGowan L, François R, Grolemund G, Hayes A, Henry L, Hester J, et al. 2019. Welcome to the Tidyverse. Journal of Open Source Software 4: 1686.

Yeaman S, Whitlock MC. 2011. The genetic architecture of adaptation under migration-selection balance: The genetic architecture of local adaptation. Evolution 65: 1897–1911.

Yeo MTS, Moyroud E. 2025. Under the rainbow: Novel insights on the mechanisms driving the development and evolution of petal pigmentation. Current Opinion in Plant Biology 86: 102743.

Yu G, Wang L-G, Han Y, He Q-Y. 2012. clusterProfiler: An R package for comparing biological themes among gene clusters. OMICS: A Journal of Integrative Biology 16: 284–287.

Yuan Y-W, Byers KJ, Bradshaw H. 2013. The genetic control of flower–pollinator specificity. Current Opinion in Plant Biology 16: 422–428.

