## Supplementary Informations for "Canalized gene regulatory networks stabilize floral polymorphism and enable modular transgressive expression"

### Supporting Information

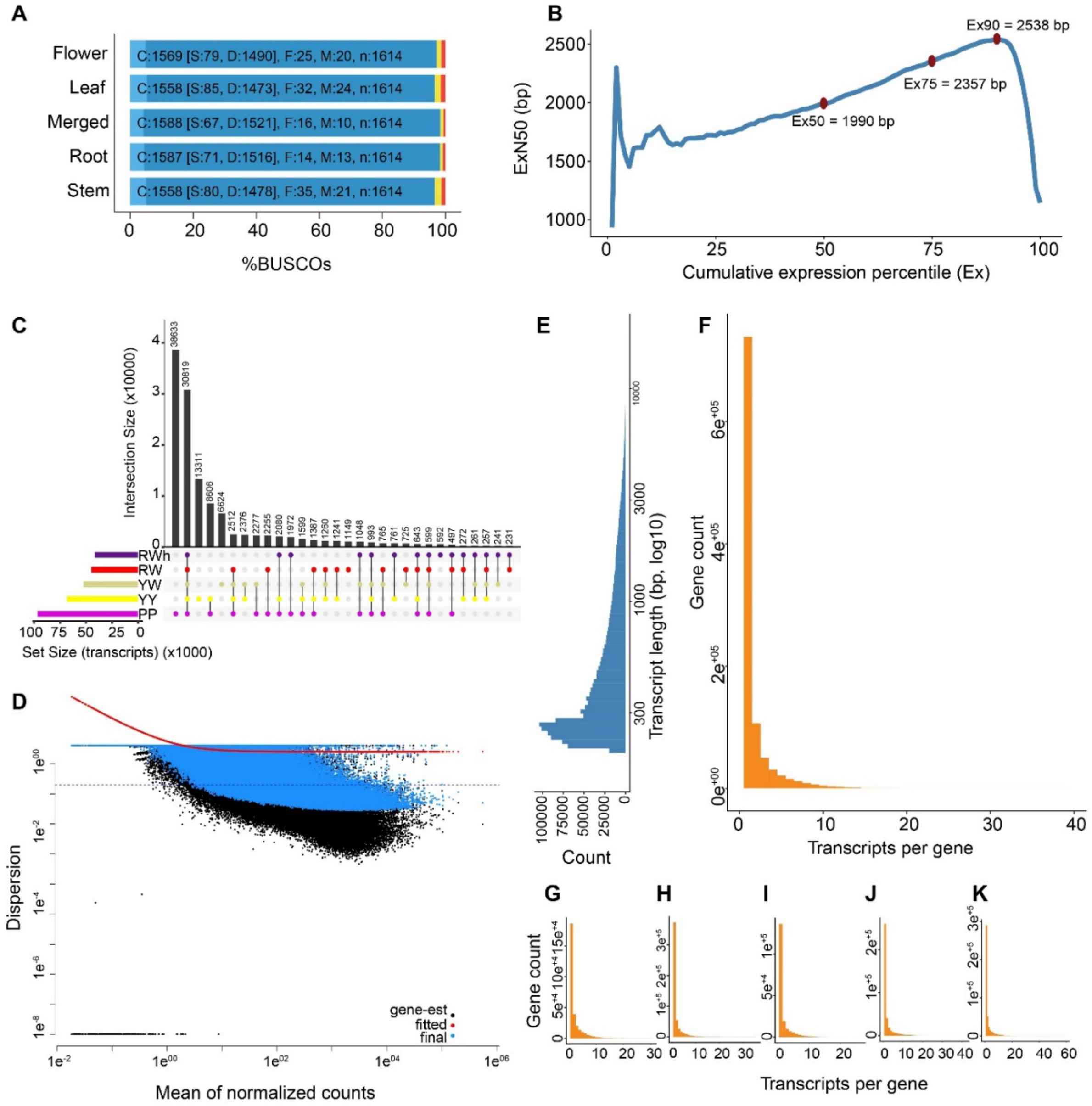


### Supplementary Fig. S1 Transcriptome assembly quality, expression characteristics, and transcript diversity.

(**A**) BUSCO completeness assessment of transcriptome assemblies generated from different tissues (flower, leaf, root, stem) and the merged assembly. Bars indicate the percentage of complete (single-copy and duplicated), fragmented, and missing BUSCOs. (**B**) ExN50 plot showing transcript contiguity as a function of cumulative expression percentile. Ex50, Ex75, and Ex90 values indicate the N50 of transcripts accounting for the top 50%, 75%, and 90% of total expression, respectively. (**C**) UpSet plot showing overlap of expressed transcripts among flower-color morphs (PP, RW, RWh, YW, YY). Bars indicate intersection sizes, and horizontal bars show total transcript counts per morph. (**D**) Mean–dispersion relationship for normalized gene counts. Black points represent genes prior to dispersion fitting, blue points represent final dispersion estimates, and the red line indicates the fitted dispersion trend. (**E**) Distribution of transcript lengths in the merged transcriptome. (**F**) Distribution of the number of transcripts per gene in the merged annotation. (**G**–**K**) Distributions of transcripts per gene for each flower-color morph (PP, RW, RWh, YW, YY), shown separately to illustrate morph-specific transcript diversity.

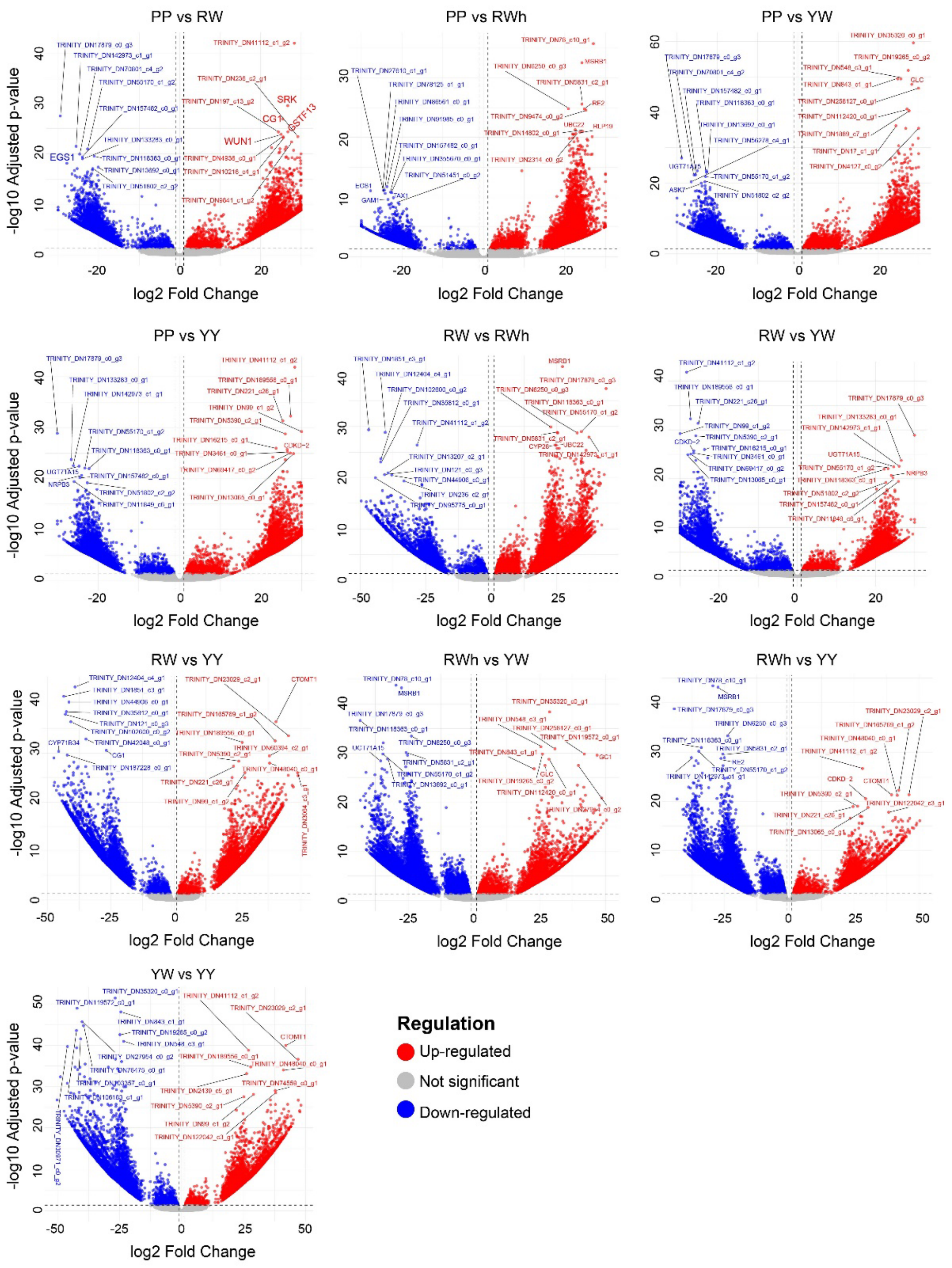


**Supplementary Fig. S2 Pairwise differential gene expression among flower-color morphs.**

Volcano plots showing differential gene expression across all pairwise comparisons among morphs (PP, RW, RWh, YW, YY). Each point represents a transcript, plotted by $\log_2$ fold change (x-axis) and $-\log_{10}$ adjusted p-value (y-axis). Fold change is calculated relative to the first morph in each comparison: negative values (blue) indicate higher expression in the first morph, whereas positive values (red) indicate higher expression in the second morph. Gray points denote non-significant transcripts. Vertical dashed lines indicate fold-change thresholds, and the horizontal dashed line indicates the significance cutoff. Selected candidate genes associated with pigment biosynthesis and regulatory pathways are labeled.

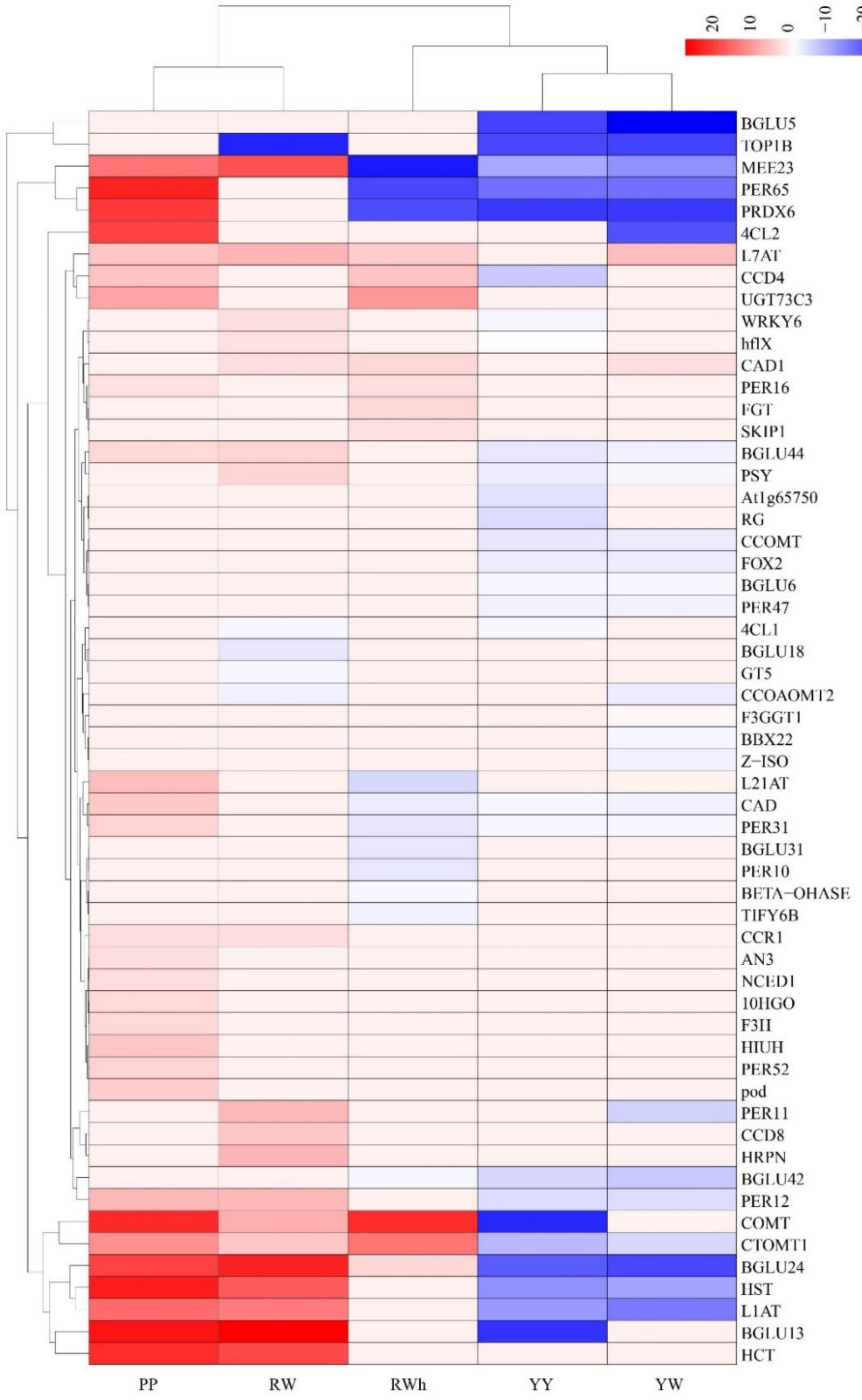


### Supplementary Fig. S3 Gene expression heatmap for pigment- and metabolism-associated genes

Hierarchically clustered heatmap showing standardized expression (Z-scores) of selected genes involved in phenylpropanoid, flavonoid, carotenoid, and stress-related pathways across floral morphs. Dendrograms indicate gene clustering based on expression similarity. The transgressive morphotype (RWh) exhibits intermediate, parent-like, and transgressive expression patterns across different gene clusters.

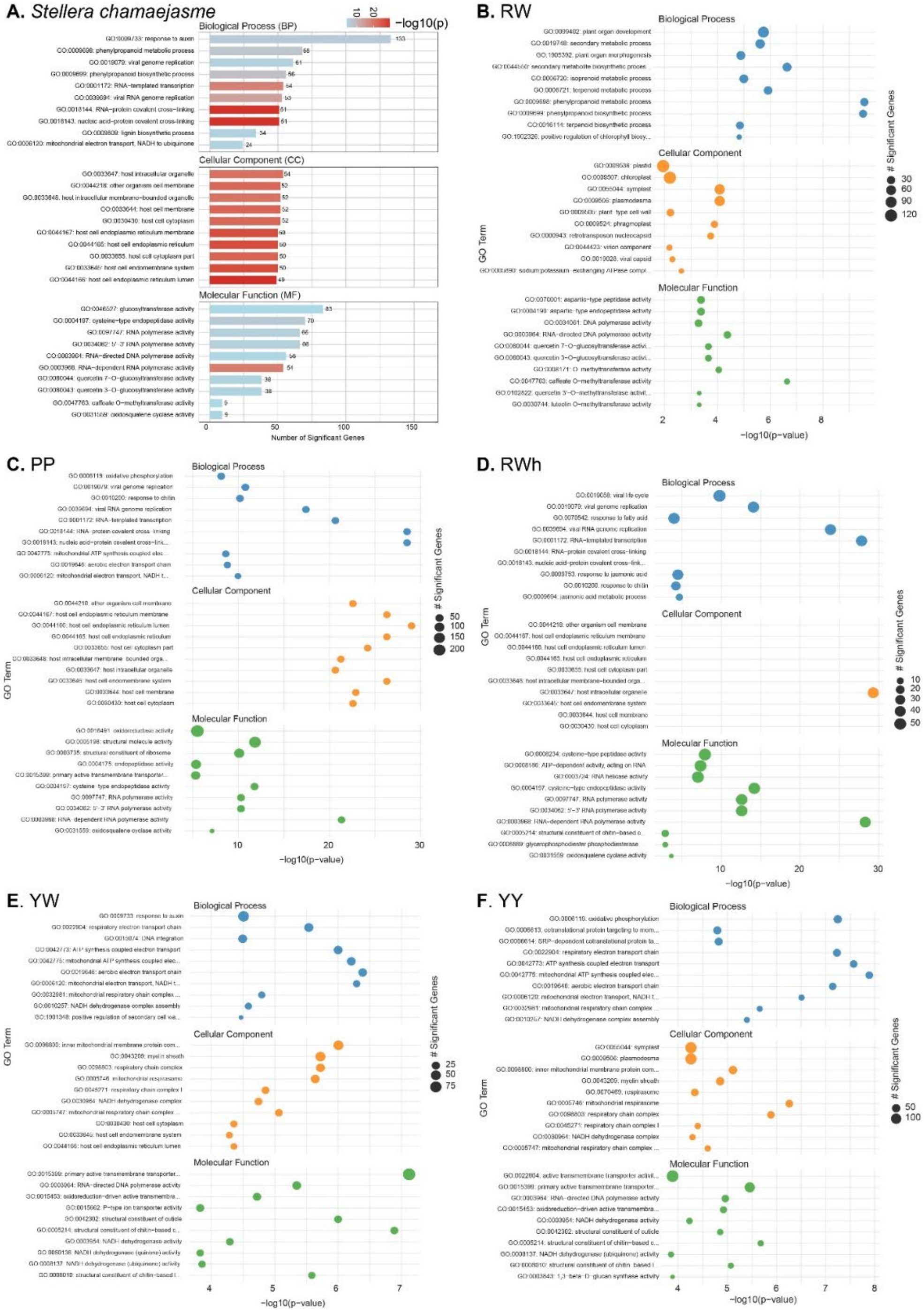


**Supplementary Fig. S4 Gene Ontology enrichment of differentially expressed genes across flower-color morphs.**

(**A**) Summary of Gene Ontology (GO) enrichment for all differentially expressed genes in *Stellera chamaejasme*, grouped by Biological Process (BP), Cellular Component (CC), and Molecular Function (MF). Bars indicate the number of significant genes per GO term, colored by $-\log_{10}(P)$ value. (**B**–**F**) GO enrichment results for differentially expressed genes specific to each flower-color morph: (**B**) red–white (RW), (**C**) pure pink (PP), (**D**) red–white transgressive (RWh), (**E**) yellow–white (YW), (**F**) pure yellow (YY). Dots represent enriched GO terms, with dot size proportional to the number of significant genes and color indicating GO category (BP, CC, MF). The x-axis shows enrichment significance as $-\log_{10}(P)$.

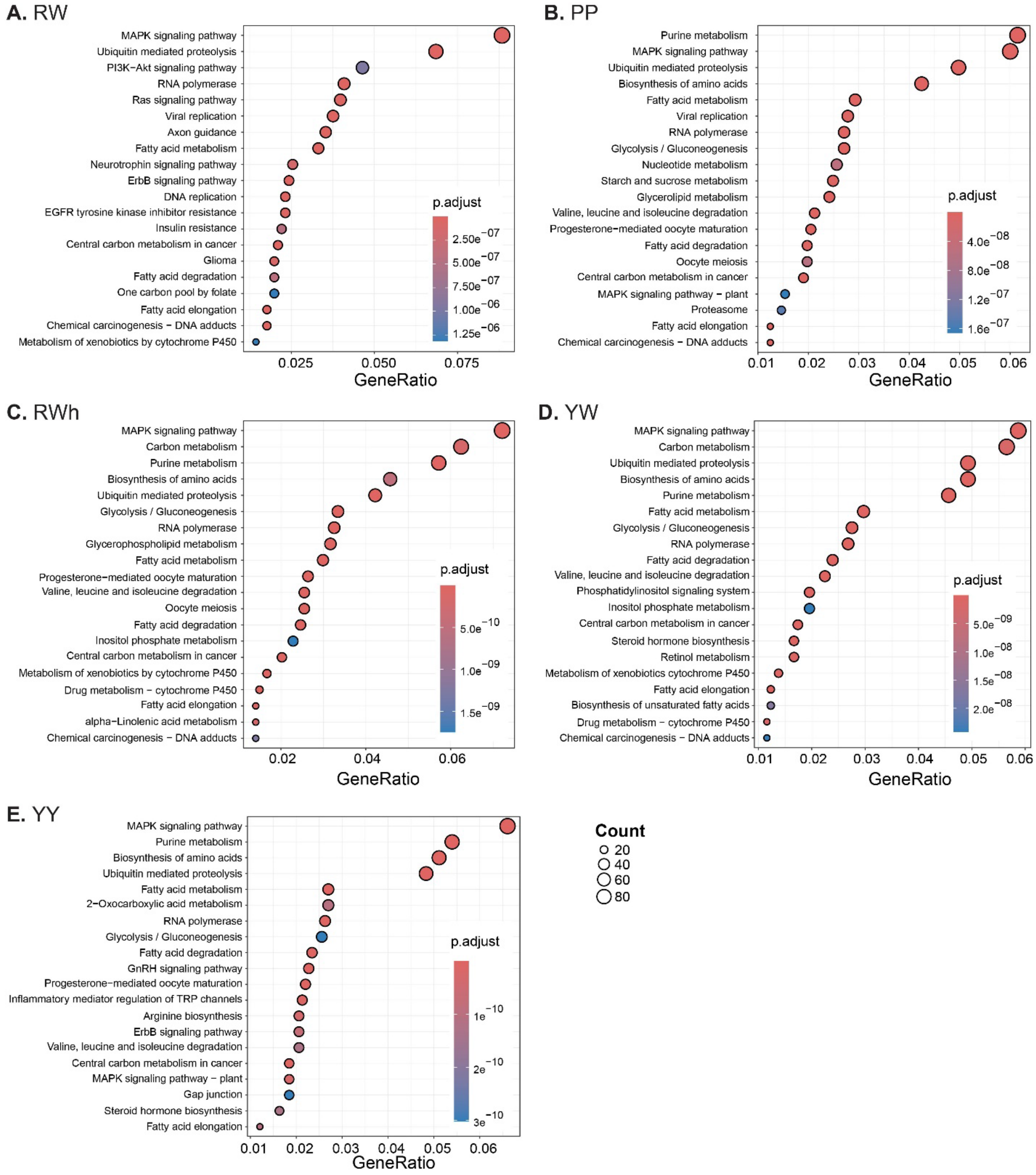


**Supplementary Fig. S5 KEGG pathway enrichment of differentially expressed genes among flower-color morphs.**

(**A**–**E**) Dot plots showing significantly enriched KEGG pathways for differentially expressed genes associated with each flower-color morph: (**A**) red–white (RW), (**B**) pure pink (PP), (**C**) red–white transgressive (RWh), (**D**) yellow–white (YW), (**E**) pure yellow (YY). Each dot represents an enriched KEGG pathway. The x-axis shows the gene ratio (proportion of genes assigned to a given pathway), dot size indicates the number of genes associated with each pathway, and color denotes the adjusted *p* (*p.adjust*). Only significantly enriched pathways are shown.

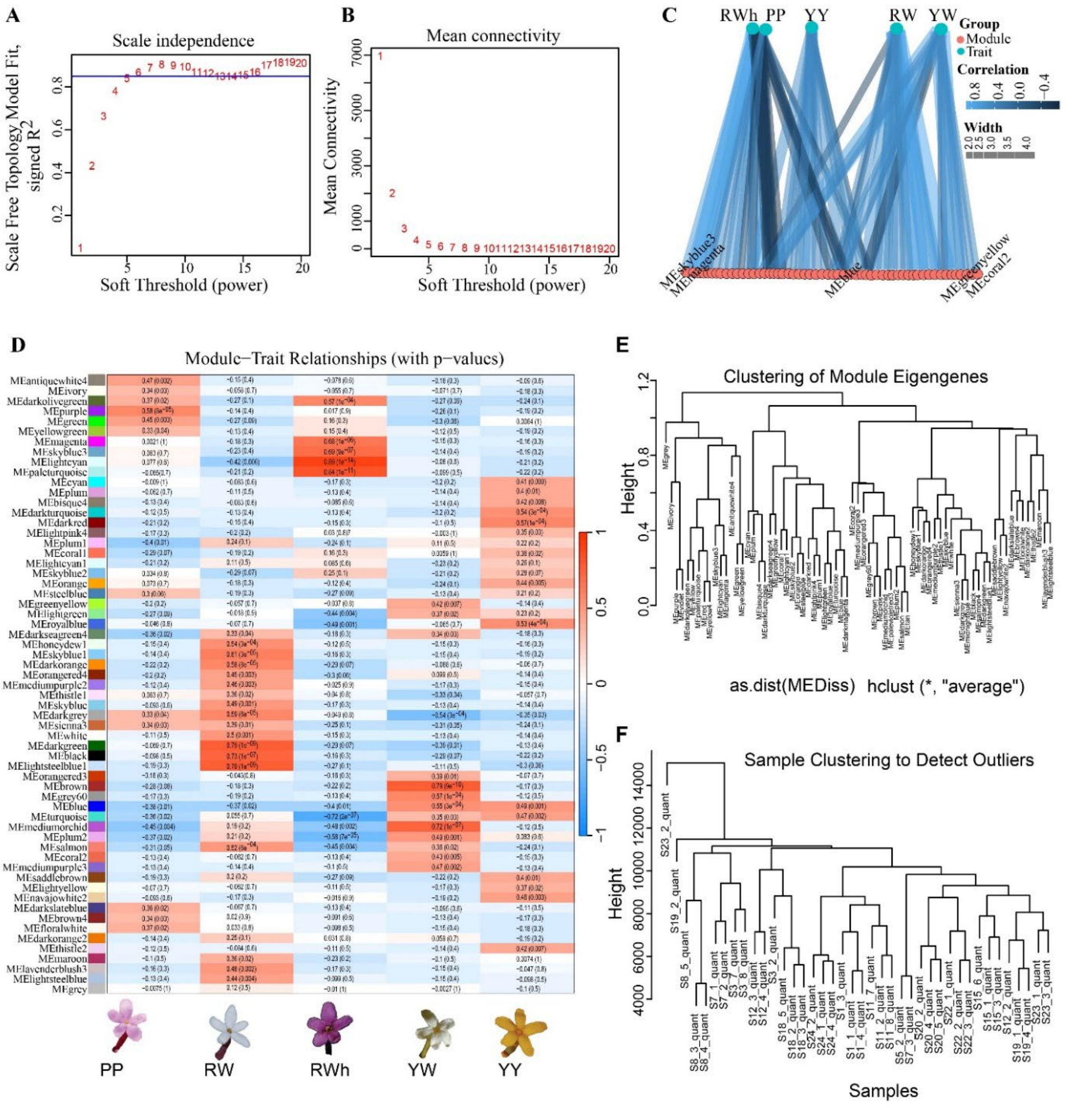


### Supplementary Fig. S6 Weighted gene co-expression network analysis (WGCNA) identifies modules associated with flower-color morphs.

(**A**) Scale-free topology model fit (signed $R^2$) as a function of soft-thresholding power. The selected power achieves approximate scale-free topology while maintaining network connectivity. (**B**) Mean connectivity across genes for each tested soft-thresholding power, illustrating the trade-off between network sparsity and connectivity. (**C**) Network visualization showing correlations between module eigengenes (modules) and flower-color morph traits (PP, RW, RWh, YW, YY). Edge width represents the strength of correlation, and color indicates correlation direction and magnitude. (**D**) Heatmap of module–trait relationships. Colors indicate Pearson correlation coefficients between module eigengenes and morph traits, with corresponding *p* shown in each cell. Modules are displayed on the y-axis and flower-color morphs on the x-axis. (**E**) Hierarchical clustering of module eigengenes based on eigengene dissimilarity, illustrating relationships among co-expression modules. (**F**) Hierarchical clustering of samples based on global gene expression profiles, used to assess sample similarity and detect potential outliers.

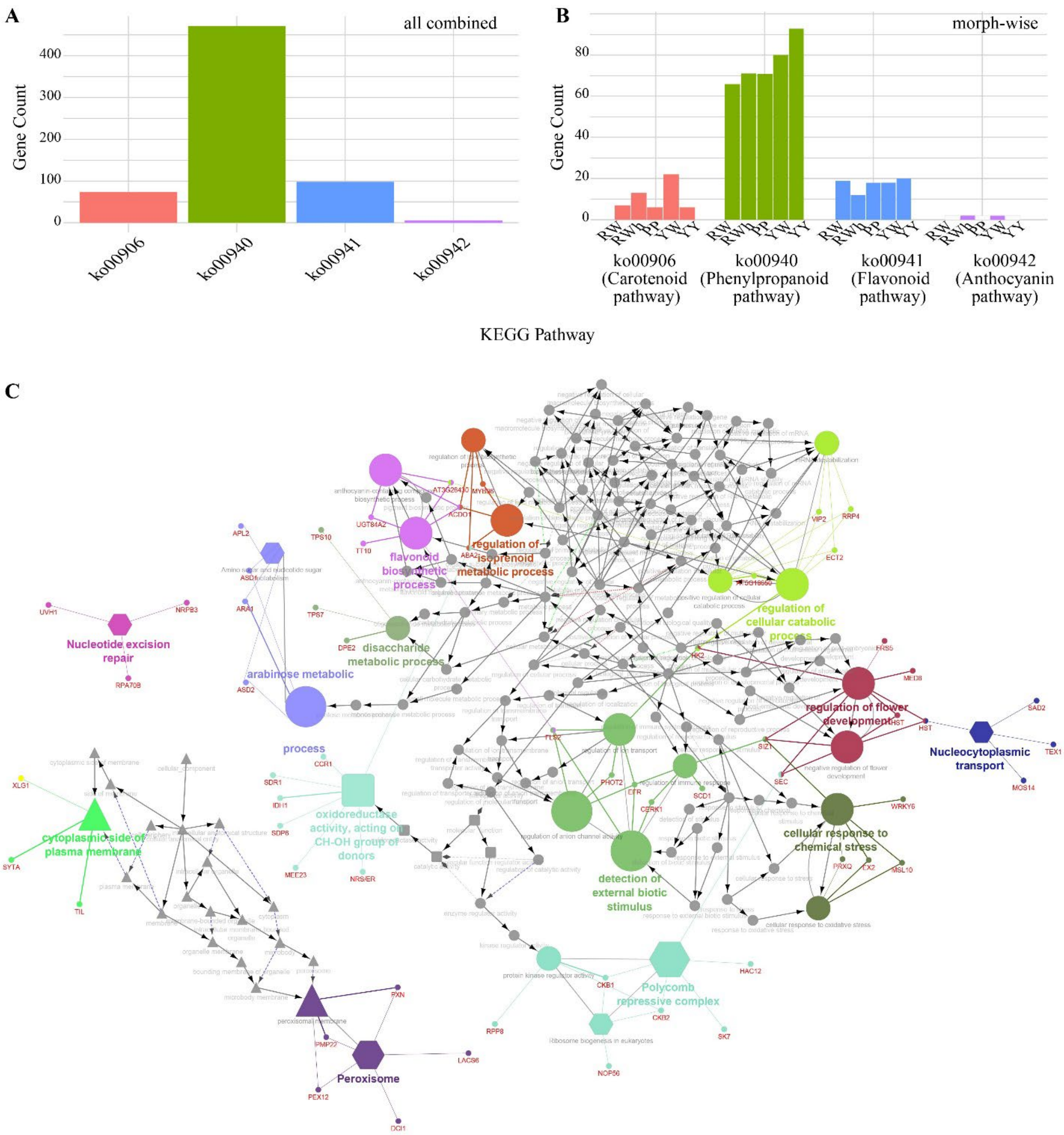


### Supplementary Fig. S7 Pathway representation and functional network structure of pigment-associated genes.

(**A**) Number of genes assigned to four pigment-related KEGG pathways (carotenoid, phenylpropanoid, flavonoid, and anthocyanin biosynthesis) across all flower-color morphs combined. (**B**) Morph-wise distribution of genes assigned to the same four pigment-related KEGG pathways. Bars indicate gene counts for each morph (PP, RW, RWh, YW, YY). (**C**) Functional enrichment network of Gene Ontology (GO) terms associated with pigment-related genes. Nodes represent enriched GO terms, with node size proportional to gene count. Edges indicate shared genes between GO terms. Colors denote functional clusters, and representative biological processes are labeled.

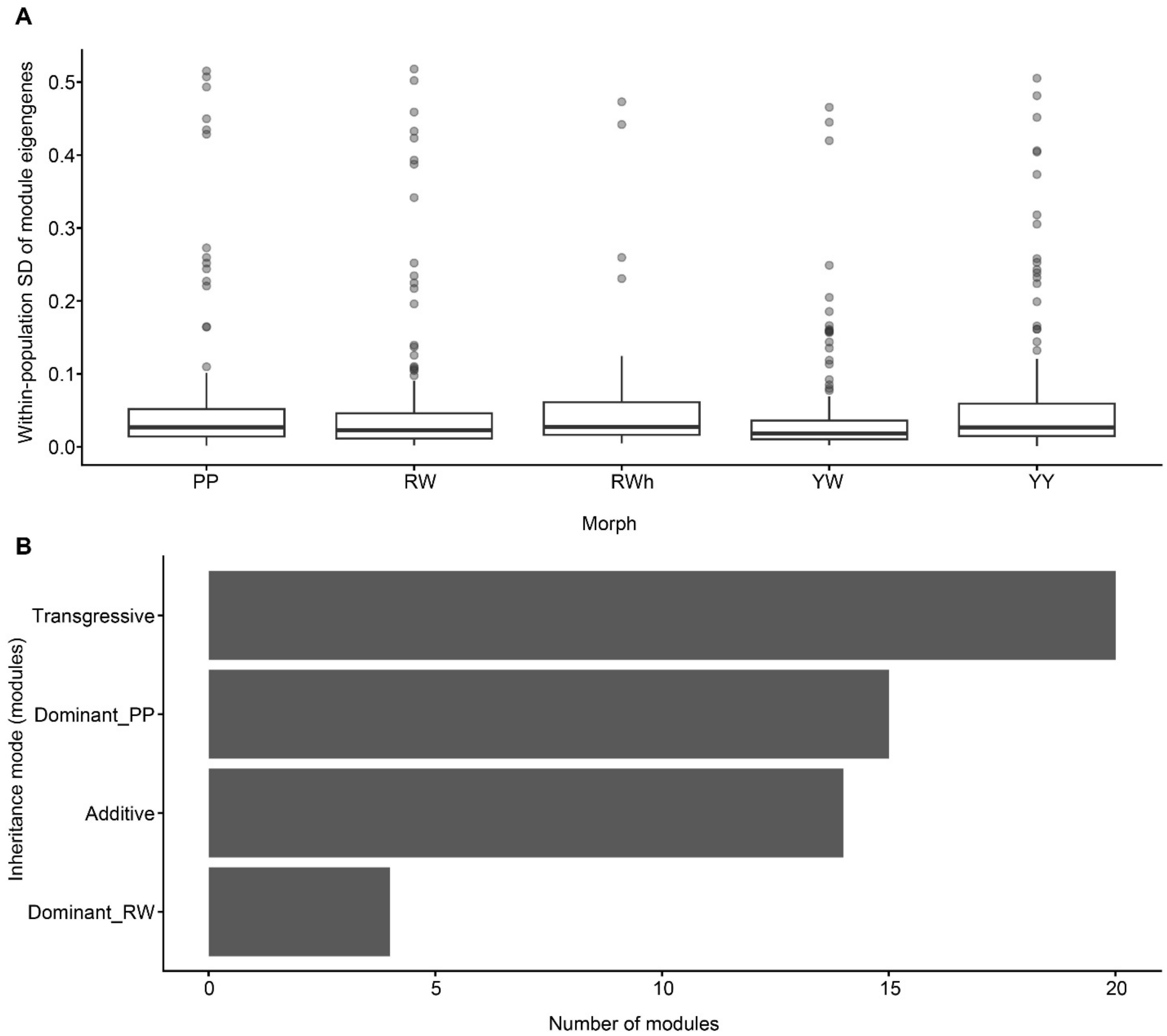


**Supplementary Fig. S8 Within-morph expression variance and inheritance patterns of co-expression modules.**

(**A**) Within-population standard deviation (SD) of module eigengene expression for each flower-color morph (PP, RW, RWh, YW, YY). Boxplots summarize the distribution of within-morph variance across co-expression modules, with points indicating individual modules. (**B**) Number of co-expression modules classified into different regulatory inheritance modes in hybrids, including additive, dominant (PP or RW), and transgressive inheritance.

### Supplementary Tables

**Supplementary Table S1** Population sampling, geographic coordinates, and experimental design.

| Morph | Population | Voucher | lat | long | Elevation (m a.s.l.) | Locality | Individuals |
|---|---|---|---|---|---|---|---|
| Pure pink morph (PP) | S3 | Deng-9525 | 30.04246 | 101.234481 | 3452.50 | Yajiang, Sichuan, China | 2, 7, 8 |
| | S8 | Deng-9533 | 30.088187 | 101.005551 | 2676.00 | Yajiang, Sichuan, China | 3, 4, 5 |
| | S23 | Deng-9813 | 30.362201 | 100.294758 | 3540.00 | Litang, Sichuan, China | 1, 2, 3 |
| Red-white morph (RW) | S15 | Deng-9629 | 29.32433 | 98.673971 | 3682.00 | Mangkang, Xizang, China | 1, 3, 6 |
| | S20 | Deng-9736 | 31.334996 | 97.214567 | 3325.50 | Changdu, Tibet, China | 1, 4, 5 |
| | S22 | Deng-9785 | 31.571412 | 100.061128 | 3356.50 | Ganzi, Sichuan, China | 1, 2, 3 |
| Red-white transgressive morph (RWh) | S5 | Deng-9527 | 30.042501 | 101.234494 | 3452.50 | Yajiang, Sichuan, China | 2 |
| | S7 | Deng-9529 | 30.042534 | 101.234446 | 3452.50 | Yajiang, Sichuan, China | 1,2 |
| Yellow-white morph (YW) | S1 | Deng-9521 | 29.996448 | 101.879447 | 3292.00 | Kangding, Sichuan, China | 1, 3, 4 |
| | S18 | Deng-9715 | 29.802221 | 98.533424 | 4210.00 | Mangkang, Xizang, China | 2, 3, 5 |
| | S24 | Deng-9822 | 29.986559 | 100.297718 | 3967.00 | Litang, Sichuan, China | 1, 2, 4 |
| Pure yellow morph (YY) | S11 | Deng-9563 | 29.173344 | 101.469558 | 3429.50 | Jiulong, Sichuan, China | 2, 7, 8 |
| | S12 | Deng-9564 | 26.541212 | 100.001661 | 3085.50 | Jianchuan, Dali, Yunnan, China | 2, 3, 4 |
| | S19 | Deng-9726 | 29.952495 | 98.380973 | 3858.50 | Mangkang, Xizang, China | 1, 2, 4 |

**Supplementary Table S2** Basic information of the population's vouchers, transcriptome sequencing, and alignment rates of RNA libraries of four tissues of four different morphs of *Stellera chamaejasme.*

| Morphs code | Voucher | Population tissues | Library | Pooled libraries | Read contents mapping rate (%) | Raw Reads | Clean Reads | Raw Base (G) | Clean Base (G) | Effective Rate (%) | Error Rate (%) | Q20 (%) | Q30 (%) | GC Content (%) |
|---|---|---|---|---|---|---|---|---|---|---|---|---|---|---|
| PP | Deng-9525 | S3-2-Flower | FRAS202235638-1r | S3_2 merged | 96.92% | 23,481,733 | 22,421,800 | 7.04 | 6.73 | 95.49 | 0.02 | 98.3 | 94.83 | 46.89 |
| | | S3-2-Leaf | FRAS202235637-1r | | | 22,239,940 | 21,301,650 | 6.67 | 6.39 | 95.78 | 0.03 | 98.18 | 94.53 | 47.25 |
| | | S3-2-Root | FRAS202235635-1r | | | 23,232,371 | 22,461,943 | 6.97 | 6.74 | 96.68 | 0.03 | 98.13 | 94.42 | 46.57 |
| | | S3-2-Stem | FRAS202235636-1r | | | 21,562,564 | 20,830,366 | 6.47 | 6.25 | 96.6 | 0.03 | 98.05 | 94.2 | 47.08 |
| | | S3-7-Flower | FRAS202235642-1r | S3_7 merged | 96.87% | 23,341,911 | 22,377,119 | 7 | 6.71 | 95.87 | 0.03 | 98.15 | 94.47 | 46.95 |
| | | S3-7-Leaf | FRAS202235641-1r | | | 23,887,900 | 22,851,715 | 7.17 | 6.86 | 95.66 | 0.03 | 98.21 | 94.59 | 47.35 |
| | | S3-7-Root | FRAS202235639-1r | | | 23,181,423 | 22,335,358 | 6.95 | 6.7 | 96.35 | 0.02 | 98.34 | 94.85 | 45.77 |
| | | S3-7-Stem | FRAS202235640-1r | | | 25,456,172 | 24,396,316 | 7.64 | 7.32 | 95.84 | 0.02 | 98.18 | 94.47 | 46.88 |
| | | S3-8-Flower | FRAS202235646-1r | S3-8 merged | 96.96% | 20,678,885 | 19,922,984 | 6.2 | 5.98 | 96.34 | 0.03 | 98.05 | 94.23 | 47.41 |
| | | S3-8-Leaf | FRAS202235645-1r | | | 22,778,686 | 22,050,038 | 6.83 | 6.62 | 96.8 | 0.02 | 98.3 | 94.82 | 47.03 |
| | | S3-8-Root | FRAS202235643-1r | | | 22,647,253 | 21,908,454 | 6.79 | 6.57 | 96.74 | 0.03 | 98.14 | 94.36 | 45.23 |
| | | S3-8-Stem | FRAS202235644-1r | | | 24,399,416 | 23,692,579 | 7.32 | 7.11 | 97.1 | 0.03 | 98.12 | 94.39 | 46.47 |
| PP | Deng-9533 | S8-3-Flower | FRAS202235650-1r | S8-3 merged | 96.92% | 20,985,311 | 20,314,151 | 6.3 | 6.09 | 96.8 | 0.03 | 98.07 | 94.33 | 47.62 |
| | | S8-3-Leaf | FRAS202235649-1r | | | 23,274,893 | 22,556,153 | 6.98 | 6.77 | 96.91 | 0.03 | 98.12 | 94.41 | 47.45 |
| | | S8-3-Root | FRAS202235647-1r | | | 22,480,264 | 21,702,029 | 6.74 | 6.51 | 96.54 | 0.03 | 97.98 | 94.08 | 46.86 |

| Morphs code | Voucher | Population tissues | Library | Pooled libraries | Read contents mapping rate (%) | Raw Reads | Clean Reads | Raw Base (G) | Clean Base (G) | Effective Rate (%) | Error Rate (%) | Q20 (%) | Q30 (%) | GC Content (%) |
|---|---|---|---|---|---|---|---|---|---|---|---|---|---|---|
| | | S8-3-Stem | FRAS202235648-1r | | | 23,287,484 | 22,457,567 | 6.99 | 6.74 | 96.44 | 0.03 | 98.13 | 94.38 | 46.45 |
| | | S8-4-Flower | FRAS202235654-1r | S8-4 merged | 96.70% | 22,101,341 | 21,334,098 | 6.63 | 6.4 | 96.53 | 0.03 | 97.84 | 93.75 | 46.75 |
| | | S8-4-Leaf | FRAS202235653-1r | | | 21,892,036 | 21,039,962 | 6.57 | 6.31 | 96.11 | 0.03 | 98.1 | 94.36 | 45.76 |
| | | S8-4-Root | FRAS202235651-1r | | | 22,091,917 | 21,372,871 | 6.63 | 6.41 | 96.75 | 0.03 | 98.06 | 94.3 | 47.05 |
| | | S8-4-Stem | FRAS202235652-1r | | | 22,083,630 | 21,302,374 | 6.63 | 6.39 | 96.46 | 0.02 | 98.24 | 94.66 | 46.5 |
| | | S8-5-Flower | FRAS202235658-1r | S8-5 merged | 96.58% | 23,796,819 | 22,867,968 | 7.14 | 6.86 | 96.1 | 0.02 | 98.14 | 94.5 | 47.3 |
| | | S8-5-Leaf | FRAS202235657-1r | | | 24,414,773 | 23,537,780 | 7.32 | 7.06 | 96.41 | 0.03 | 98.07 | 94.3 | 47 |
| | | S8-5-Root | FRAS202235655-1r | | | 22,138,565 | 21,324,276 | 6.64 | 6.4 | 96.32 | 0.03 | 98.03 | 94.16 | 42.36 |
| | | S8-5-Stem | FRAS202235656-1r | | | 21,292,938 | 20,651,607 | 6.39 | 6.2 | 96.99 | 0.02 | 98.28 | 94.72 | 46.7 |
| PP | Deng-9813 | S23-1-Flower | FRAS202235662-1r | S23-1 merged | 96.04% | 21,954,122 | 21,128,816 | 6.59 | 6.34 | 96.24 | 0.03 | 98.01 | 94.14 | 47.04 |
| | | S23-1-Leaf | FRAS202235661-1r | | | 22,375,665 | 21,583,527 | 6.71 | 6.48 | 96.46 | 0.02 | 98.26 | 94.74 | 46.86 |
| | | S23-1-Root | FRAS202235659-1r | | | 21,646,842 | 20,848,588 | 6.49 | 6.25 | 96.31 | 0.03 | 98.11 | 94.31 | 45.89 |
| | | S23-1-Stem | FRAS202235660-1r | | | 20,838,598 | 20,126,854 | 6.25 | 6.04 | 96.58 | 0.03 | 98.06 | 94.29 | 46.16 |
| | | S23-2-Flower | FRAS202235666-1r | S23-2 merged | 95.94% | 21,542,597 | 20,744,507 | 6.46 | 6.22 | 96.3 | 0.03 | 98.17 | 94.54 | 46.25 |
| | | S23-2-Leaf | FRAS202235665-1r | | | 20,738,263 | 19,993,448 | 6.22 | 6 | 96.41 | 0.02 | 98.2 | 94.57 | 45.7 |
| | | S23-2-Root | FRAS202235663-1r | | | 23,084,198 | 22,303,721 | 6.93 | 6.69 | 96.62 | 0.02 | 98.22 | 94.63 | 46.25 |
| | | S23-2-Stem | FRAS202235664-1r | | | 20,286,007 | 19,508,778 | 6.09 | 5.85 | 96.17 | 0.03 | 98.12 | 94.4 | 46.08 |

| Morphs code | Voucher | Population tissues | Library | Pooled libraries | Read contents mapping rate (%) | Raw Reads | Clean Reads | Raw Base (G) | Clean Base (G) | Effective Rate (%) | Error Rate (%) | Q20 (%) | Q30 (%) | GC Content (%) |
|---|---|---|---|---|---|---|---|---|---|---|---|---|---|---|
| | | S23-3-Flower | FRAS202235670-1r | S23-3 merged | 96.23% | 19,854,020 | 19,142,724 | 5.96 | 5.74 | 96.42 | 0.03 | 98.13 | 94.36 | 46.52 |
| | | S23-3-Leaf | FRAS202235669-1r | | | 22,503,976 | 21,663,099 | 6.75 | 6.5 | 96.26 | 0.02 | 98.25 | 94.64 | 45.85 |
| | | S23-3-Root | FRAS202235667-1r | | | 22,682,103 | 21,899,812 | 6.8 | 6.57 | 96.55 | 0.03 | 98.14 | 94.46 | 46.08 |
| | | S23-3-Stem | FRAS202235668-1r | | | 22,235,054 | 21,363,106 | 6.67 | 6.41 | 96.08 | 0.03 | 97.95 | 93.97 | 45.73 |
| RW | Deng-9629 | S15-1-Flower | FRAS202235674-1r | S15-1 merged | 96.16% | 20,585,523 | 19,853,700 | 6.18 | 5.96 | 96.44 | 0.03 | 97.97 | 94.06 | 46.62 |
| | | S15-1-Leaf | FRAS202235673-1r | | | 21,807,777 | 21,108,399 | 6.54 | 6.33 | 96.79 | 0.02 | 98.25 | 94.72 | 46.8 |
| | | S15-1-Root | FRAS202235671-1r | | | 21,168,765 | 20,360,802 | 6.35 | 6.11 | 96.18 | 0.03 | 98.15 | 94.5 | 46.36 |
| | | S15-1-Stem | FRAS202235672-1r | | | 20,955,479 | 20,288,556 | 6.29 | 6.09 | 96.82 | 0.03 | 98.12 | 94.38 | 46.28 |
| | | S15-3-Flower | FRAS202235678-1r | S15-3 merged | 95.87% | 22,818,652 | 22,050,466 | 6.85 | 6.62 | 96.63 | 0.03 | 98.06 | 94.25 | 46.94 |
| | | S15-3-Leaf | FRAS202235677-1r | | | 19,967,539 | 19,302,370 | 5.99 | 5.79 | 96.67 | 0.03 | 97.94 | 93.96 | 47.28 |
| | | S15-3-Root | FRAS202235675-1r | | | 23,023,623 | 22,345,682 | 6.91 | 6.7 | 97.06 | 0.03 | 98.06 | 94.22 | 45.86 |
| | | S15-3-Stem | FRAS202235676-1r | | | 22,849,191 | 22,044,995 | 6.85 | 6.61 | 96.48 | 0.03 | 98.03 | 94.19 | 46.64 |
| | | S15-6-Flower | FRAS202235682-1r | S15-6 merged | 96.12% | 21,888,319 | 21,139,812 | 6.57 | 6.34 | 96.58 | 0.03 | 98.06 | 94.28 | 47.32 |
| | | S15-6-Leaf | FRAS202235681-1r | | | 21,888,194 | 21,126,437 | 6.57 | 6.34 | 96.52 | 0.02 | 98.19 | 94.59 | 47.06 |
| | | S15-6-Root | FRAS202235679-1r | | | 22,934,583 | 22,224,689 | 6.88 | 6.67 | 96.9 | 0.03 | 97.94 | 93.98 | 46.22 |
| | | S15-6-Stem | FRAS202235680-1r | | | 24,703,027 | 23,923,716 | 7.41 | 7.18 | 96.85 | 0.03 | 98.05 | 94.23 | 46.5 |
| RW | Deng-9736 | S20-2-leaf | FRAS202236054-1r | S20-2 merged | 97.07% | 33,661,218 | 32,135,643 | 10.1 | 9.64 | 95.47 | 0.02 | 98.6 | 95.66 | 47.18 |

| Morphs code | Voucher | Population tissues | Library | Pooled libraries | Read contents mapping rate (%) | Raw Reads | Clean Reads | Raw Base (G) | Clean Base (G) | Effective Rate (%) | Error Rate (%) | Q20 (%) | Q30 (%) | GC Content (%) |
|---|---|---|---|---|---|---|---|---|---|---|---|---|---|---|
| | | S20-2-root | FRAS202236052-1r | | | 24,099,886 | 23,088,355 | 7.23 | 6.93 | 95.8 | 0.02 | 98.48 | 95.34 | 46.55 |
| | | S20-2-root | FRAS202236055-1r | | | 22,947,354 | 21,827,789 | 6.88 | 6.55 | 95.12 | 0.02 | 98.58 | 95.56 | 47.3 |
| | | S20-2-stem | FRAS202236053-1r | | | 24,493,709 | 23,655,341 | 7.35 | 7.1 | 96.58 | 0.02 | 98.3 | 94.91 | 46.47 |
| | | S20-4-Flower | FRAS202235690-1r | S20-4 merged | 96.40% | 20,558,285 | 19,911,407 | 6.17 | 5.97 | 96.85 | 0.03 | 98.07 | 94.32 | 47.64 |
| | | S20-4-Leaf | FRAS202235689-1r | | | 21,627,393 | 20,942,030 | 6.49 | 6.28 | 96.83 | 0.03 | 98.06 | 94.31 | 47.05 |
| | | S20-4-Root | FRAS202235687-1r | | | 21,789,943 | 21,036,654 | 6.54 | 6.31 | 96.54 | 0.03 | 98.11 | 94.41 | 46.36 |
| | | S20-4-Stem | FRAS202235688-1r | | | 20,317,329 | 19,716,151 | 6.1 | 5.91 | 97.04 | 0.03 | 98.1 | 94.33 | 46.5 |
| | | S20-5-Flower | FRAS202235694-1r | S20-5 merged | 96.26% | 19,856,032 | 19,121,852 | 5.96 | 5.74 | 96.3 | 0.03 | 98.15 | 94.48 | 46.94 |
| | | S20-5-Leaf | FRAS202235693-2a | | | 22,798,374 | 21,961,019 | 6.84 | 6.59 | 96.33 | 0.03 | 97.62 | 93.47 | 47.34 |
| | | S20-5-Root | FRAS202235691-1r | | | 20,500,971 | 19,847,056 | 6.15 | 5.95 | 96.81 | 0.03 | 98.24 | 94.62 | 46.45 |
| | | S20-5-Stem | FRAS202235692-1r | | | 21,460,046 | 20,737,676 | 6.44 | 6.22 | 96.63 | 0.02 | 98.32 | 94.86 | 47.28 |
| RW | Deng-9785 | S22-1-Flower | FRAS202235698-1r | S22-1 merged | 96.91% | 20,515,392 | 19,760,430 | 6.15 | 5.93 | 96.32 | 0.03 | 98 | 94.17 | 46.91 |
| | | S22-1-Leaf | FRAS202235697-1r | | | 21,840,191 | 20,426,053 | 6.55 | 6.13 | 93.53 | 0.03 | 97.84 | 93.81 | 46.91 |
| | | S22-1-Root | FRAS202235695-1r | | | 20,958,287 | 20,081,643 | 6.29 | 6.02 | 95.82 | 0.03 | 98 | 94.16 | 46.53 |
| | | S22-1-Stem | FRAS202235696-1r | | | 23,979,103 | 23,051,542 | 7.19 | 6.92 | 96.13 | 0.02 | 98.16 | 94.51 | 46.45 |
| | | S22-2-Flower | FRAS202235702-1r | S22-2 merged | 96.68% | 21,120,546 | 20,235,134 | 6.34 | 6.07 | 95.81 | 0.03 | 98.09 | 94.34 | 46.34 |
| | | S22-2-Leaf | FRAS202235701-1r | | | 21,847,136 | 20,916,989 | 6.55 | 6.28 | 95.74 | 0.03 | 98.08 | 94.31 | 46.47 |

| Morphs code | Voucher | Population tissues | Library | Pooled libraries | Read contents mapping rate (%) | Raw Reads | Clean Reads | Raw Base (G) | Clean Base (G) | Effective Rate (%) | Error Rate (%) | Q20 (%) | Q30 (%) | GC Content (%) |
|---|---|---|---|---|---|---|---|---|---|---|---|---|---|---|
| | | S22-2-Root | FRAS202235699-1r | | | 21,563,047 | 20,770,260 | 6.47 | 6.23 | 96.32 | 0.02 | 98.16 | 94.49 | 46.46 |
| | | S22-2-Stem | FRAS202235700-1r | | | 20,257,019 | 19,487,351 | 6.08 | 5.85 | 96.2 | 0.03 | 98.03 | 94.2 | 46.49 |
| | | S22-3-Flower | FRAS202235706-1r | S22-3 merged | 96.64% | 24,036,119 | 22,894,763 | 7.21 | 6.87 | 95.25 | 0.03 | 98.07 | 94.31 | 45.97 |
| | | S22-3-Leaf | FRAS202235705-1r | | | 22,424,231 | 21,346,352 | 6.73 | 6.4 | 95.19 | 0.03 | 98.07 | 94.39 | 46.12 |
| | | S22-3-Root | FRAS202235703-1r | | | 23,508,066 | 22,505,761 | 7.05 | 6.75 | 95.74 | 0.03 | 98.05 | 94.32 | 46.33 |
| | | S22-3-Stem | FRAS202235704-1r | | | 19,808,408 | 18,843,360 | 5.94 | 5.65 | 95.13 | 0.03 | 98.05 | 94.24 | 45.52 |
| RWh | Deng-9527 | S5-2-Flower | FRAS202235718-1r | S5-2 merged | 96.90% | 23,449,933 | 22,544,609 | 7 | 7 | 96.14 | 0.03 | 97.67 | 93.42 | 45.86 |
| | | S5-2-Leaf | FRAS202235717-1r | | | 21,161,527 | 20,346,387 | 6 | 6 | 96.15 | 0.03 | 98 | 94.1 | 45.19 |
| | | S5-2-Root | FRAS202235715-1r | | | 19,854,329 | 19,263,654 | 5.96 | 5.78 | 97.02 | 0.03 | 98.05 | 94.15 | 44.71 |
| | Deng-9529 | S7-1-Flower | FRAS202235710-1r | S7-1 merged | 96.20% | 21,144,844 | 20,384,895 | 6 | 6 | 96.41 | 0.03 | 98.02 | 94.21 | 46.8 |
| | | S7-1-Leaf | FRAS202235709-1r | | | 21,616,102 | 20,590,666 | 6 | 6 | 95.26 | 0.02 | 98.32 | 94.81 | 46.22 |
| | | S7-1-Root | FRAS202235707-1b | | | 32,460,167 | 31,163,355 | 10 | 9 | 96 | 0.03 | 97.98 | 94.27 | 45.86 |
| | | S7-1-Stem | FRAS202235708-1r | | | 21,516,347 | 20,680,643 | 6 | 6 | 96.12 | 0.03 | 98.12 | 94.47 | 46.71 |
| | | S7-2-Flower | FRAS202235714-1r | S7-2 merged | 96.41% | 21,236,697 | 20,410,831 | 6 | 6 | 96.11 | 0.03 | 98.1 | 94.37 | 46.55 |
| | | S7-2-Leaf | FRAS202235713-1r | | | 27,484,118 | 26,315,823 | 8 | 8 | 95.75 | 0.03 | 98.09 | 94.37 | 46.18 |
| | | S7-2-Root | FRAS202235711-1r | | | 20,322,288 | 19,664,866 | 6 | 6 | 96.77 | 0.03 | 98.12 | 94.4 | 46.16 |
| | | S7-2-Stem | FRAS202235712-1r | | | 23,346,569 | 22,471,911 | 7 | 7 | 96.25 | 0.03 | 98.06 | 94.32 | 46.3 |

| Morphs code | Voucher | Population tissues | Library | Pooled libraries | Read contents mapping rate (%) | Raw Reads | Clean Reads | Raw Base (G) | Clean Base (G) | Effective Rate (%) | Error Rate (%) | Q20 (%) | Q30 (%) | GC Content (%) |
|---|---|---|---|---|---|---|---|---|---|---|---|---|---|---|
| YW | Deng-9521 | S1-1_Flower | FRAS202235722-1r | S1-1 merged | 96.66% | 20,740,712 | 20,084,775 | 6.22 | 6.03 | 96.84 | 0.03 | 98.05 | 94.25 | 46.74 |
| | | S1-1_Leaf | FRAS202235721-1r | | | 26,208,087 | 25,302,941 | 7.86 | 7.59 | 96.55 | 0.03 | 98.17 | 94.52 | 47.41 |
| | | S1-1_Root | FRAS202235719-1r | | | 20,309,881 | 19,619,635 | 6.09 | 5.89 | 96.6 | 0.03 | 98 | 94.14 | 45.76 |
| | | S1-1_Stem | FRAS202235720-1r | | | 22,517,264 | 21,729,321 | 6.76 | 6.52 | 96.5 | 0.03 | 98.16 | 94.52 | 46.9 |
| | | S1-3_Flower | FRAS202235726-1r | S1-3 merged | 96.44% | 22,587,927 | 21,874,569 | 6.78 | 6.56 | 96.84 | 0.02 | 98.24 | 94.56 | 46.11 |
| | | S1-3_Leaf | FRAS202235725-1r | | | 21,055,828 | 20,376,232 | 6.32 | 6.11 | 96.77 | 0.03 | 98.09 | 94.38 | 46.8 |
| | | S1-3_Root | FRAS202235723-1r | | | 24,202,400 | 23,417,277 | 7.26 | 7.03 | 96.76 | 0.03 | 98.03 | 94.19 | 45.95 |
| | | S1-3_Stem | FRAS202235724-1r | | | 21,532,268 | 20,767,214 | 6.46 | 6.23 | 96.45 | 0.03 | 98.07 | 94.32 | 46.34 |
| | | S1-4_Flower | FRAS202235730-1r | S1-4 merged | 96.72% | 22,072,666 | 21,308,812 | 6.62 | 6.39 | 96.54 | 0.03 | 98.03 | 94.22 | 46.57 |
| | | S1-4_Leaf | FRAS202235729-1r | | | 25,124,057 | 24,304,571 | 7.54 | 7.29 | 96.74 | 0.03 | 98.15 | 94.47 | 47.48 |
| | | S1-4_Root | FRAS202235727-1r | | | 23,089,988 | 22,265,443 | 6.93 | 6.68 | 96.43 | 0.02 | 98.18 | 94.53 | 46.12 |
| | | S1-4_Stem | FRAS202235728-1r | | | 23,617,591 | 22,760,948 | 7.09 | 6.83 | 96.37 | 0.03 | 98.03 | 94.24 | 47.17 |
| YW | Deng-9715 | S18-2_Flower | FRAS202235734-1r | S18-2 merged | 96.99% | 22,459,677 | 21,481,182 | 6.74 | 6.44 | 95.64 | 0.03 | 98.08 | 94.34 | 46.32 |
| | | S18-2_Leaf | FRAS202235733-1r | | | 22,405,005 | 21,489,319 | 6.72 | 6.45 | 95.91 | 0.03 | 98.08 | 94.32 | 46.24 |
| | | S18-2_Root | FRAS202235731-1r | | | 23,502,281 | 22,499,765 | 7.05 | 6.75 | 95.73 | 0.02 | 98.22 | 94.62 | 46.14 |
| | | S18-2_Stem | FRAS202235732-1r | | | 22,190,326 | 21,311,709 | 6.66 | 6.39 | 96.04 | 0.03 | 98.18 | 94.53 | 46.36 |
| | | S18-3_Flower | FRAS202235738-1r | S18-3 merged | 96.50% | 20,243,248 | 19,579,172 | 6.07 | 5.87 | 96.72 | 0.03 | 98.06 | 94.3 | 46.62 |

| Morphs code | Voucher | Population tissues | Library | Pooled libraries | Read contents mapping rate (%) | Raw Reads | Clean Reads | Raw Base (G) | Clean Base (G) | Effective Rate (%) | Error Rate (%) | Q20 (%) | Q30 (%) | GC Content (%) |
|---|---|---|---|---|---|---|---|---|---|---|---|---|---|---|
| | | S18-3_Leaf | FRAS202235737-1r | | | 21,722,729 | 21,109,082 | 6.52 | 6.33 | 97.18 | 0.02 | 98.38 | 94.9 | 46.57 |
| | | S18-3_Root | FRAS202235735-1r | | | 20,978,142 | 20,319,598 | 6.29 | 6.1 | 96.86 | 0.03 | 98.09 | 94.35 | 45.77 |
| | | S18-3_Stem | FRAS202235736-1r | | | 21,644,842 | 20,931,879 | 6.49 | 6.28 | 96.71 | 0.03 | 98 | 94.15 | 46.31 |
| | | S18-5_Flower | FRAS202235742-1r | S18-5 merged | 96.61% | 21,202,528 | 20,523,939 | 6.36 | 6.16 | 96.8 | 0.03 | 97.81 | 93.75 | 46.62 |
| | | S18-5_Leaf | FRAS202235741-1r | | | 22,495,311 | 21,644,978 | 6.75 | 6.49 | 96.22 | 0.02 | 98.23 | 94.71 | 46.89 |
| | | S18-5_Root | FRAS202235739-1r | | | 23,973,770 | 23,141,547 | 7.19 | 6.94 | 96.53 | 0.03 | 97.96 | 94.04 | 45.72 |
| | | S18-5_Stem | FRAS202235740-1r | | | 22,765,486 | 21,989,304 | 6.83 | 6.6 | 96.59 | 0.03 | 98.08 | 94.31 | 46.11 |
| YW | Deng-9822 | S24-1_Flower | FRAS202235746-1r | S24-1 merged | 96.89% | 22,194,544 | 21,457,600 | 6.66 | 6.44 | 96.68 | 0.02 | 98.26 | 94.69 | 46.3 |
| | | S24-1_Leaf | FRAS202235745-1r | | | 20,831,041 | 20,163,479 | 6.25 | 6.05 | 96.8 | 0.03 | 98.09 | 94.34 | 46.86 |
| | | S24-1_Root | FRAS202235743-1r | | | 23,059,064 | 22,229,624 | 6.92 | 6.67 | 96.4 | 0.02 | 98.18 | 94.56 | 46.3 |
| | | S24-1_Stem | FRAS202235744-1r | | | 23,018,133 | 22,326,883 | 6.91 | 6.7 | 97 | 0.03 | 98.06 | 94.33 | 46.39 |
| | | S24-2_Flower | FRAS202235750-1r | S24-2 merged | 96.77% | 21,840,193 | 21,126,763 | 6.55 | 6.34 | 96.73 | 0.03 | 98.2 | 94.59 | 46.21 |
| | | S24-2_Leaf | FRAS202235749-1r | | | 19,727,066 | 19,022,844 | 5.92 | 5.71 | 96.43 | 0.03 | 98.09 | 94.4 | 47.01 |
| | | S24-2_Root | FRAS202235747-1r | | | 20,966,881 | 20,227,979 | 6.29 | 6.07 | 96.48 | 0.03 | 98.09 | 94.37 | 46.55 |
| | | S24-2_Stem | FRAS202235748-1r | | | 22,432,640 | 21,694,771 | 6.73 | 6.51 | 96.71 | 0.03 | 97.88 | 93.87 | 46.5 |
| | | S24-4_Flower | FRAS202235754-1r | S24-4 merged | 96.30% | 21,758,630 | 20,974,073 | 6.53 | 6.29 | 96.39 | 0.03 | 98.11 | 94.43 | 46.27 |
| | | S24-4_Leaf | FRAS202235753-1r | | | 21,447,349 | 20,706,357 | 6.43 | 6.21 | 96.55 | 0.03 | 98.02 | 94.21 | 46.72 |

| Morphs code | Voucher | Population tissues | Library | Pooled libraries | Read contents mapping rate (%) | Raw Reads | Clean Reads | Raw Base (G) | Clean Base (G) | Effective Rate (%) | Error Rate (%) | Q20 (%) | Q30 (%) | GC Content (%) |
|---|---|---|---|---|---|---|---|---|---|---|---|---|---|---|
| | | S24-4_Root | FRAS202235751-1r | | | 22,361,079 | 21,603,327 | 6.71 | 6.48 | 96.61 | 0.03 | 98.07 | 94.21 | 46.13 |
| | | S24-4_Stem | FRAS202235752-1r | | | 23,216,942 | 22,323,032 | 6.97 | 6.7 | 96.15 | 0.03 | 98.1 | 94.38 | 45.84 |
| YY | Deng-9563 | S11-2_Flower | FRAS202235758-1r | S11-2 merged | 96.85% | 20,923,294 | 20,211,029 | 6.28 | 6.06 | 96.6 | 0.03 | 98.16 | 94.5 | 46.97 |
| | | S11-2_Leaf | FRAS202235757-1r | | | 21,548,088 | 20,781,114 | 6.46 | 6.23 | 96.44 | 0.02 | 98.22 | 94.69 | 47.48 |
| | | S11-2_Root | FRAS202235755-1r | | | 22,195,116 | 21,488,906 | 6.66 | 6.45 | 96.82 | 0.03 | 98.14 | 94.45 | 46.52 |
| | | S11-2_Stem | FRAS202235756-1r | | | 23,058,930 | 22,213,518 | 6.92 | 6.66 | 96.33 | 0.02 | 98.17 | 94.58 | 46.87 |
| | | S11-7_Flower | FRAS202235762-1r | S11-7 merged | 96.69% | 23,228,906 | 22,434,517 | 6.97 | 6.73 | 96.58 | 0.03 | 98.12 | 94.42 | 46.53 |
| | | S11-7_Leaf | FRAS202235761-1r | | | 20,523,253 | 19,929,961 | 6.16 | 5.98 | 97.11 | 0.03 | 97.97 | 93.98 | 47.39 |
| | | S11-7_Root | FRAS202235759-1r | | | 19,921,030 | 19,210,327 | 5.98 | 5.76 | 96.43 | 0.03 | 98.08 | 94.3 | 46.08 |
| | | S11-7_Stem | FRAS202235760-1r | | | 20,508,023 | 19,923,043 | 6.15 | 5.98 | 97.15 | 0.03 | 98.02 | 94.21 | 46.91 |
| | | S11-8_Flower | FRAS202235766-1r | S11-8 merged | 96.75% | 22,308,363 | 21,568,287 | 6.69 | 6.47 | 96.68 | 0.02 | 98.18 | 94.58 | 47.14 |
| | | S11-8_Leaf | FRAS202235765-1b | | | 32,385,387 | 31,315,921 | 9.72 | 9.39 | 96.7 | 0.03 | 98.06 | 94.53 | 47.8 |
| | | S11-8_Root | FRAS202235763-1r | | | 23,517,537 | 22,781,759 | 7.06 | 6.83 | 96.87 | 0.03 | 98.03 | 94.26 | 46.31 |
| | | S11-8_Stem | FRAS202235764-1r | | | 22,544,815 | 21,829,191 | 6.76 | 6.55 | 96.83 | 0.03 | 98.18 | 94.55 | 47.06 |
| YY | Deng-9564 | S12-2_Flower | FRAS202235770-1r | S12-2 merged | 96.20% | 22,341,767 | 21,640,703 | 6.7 | 6.49 | 96.86 | 0.03 | 98.01 | 94.19 | 46.78 |
| | | S12-2_Leaf | FRAS202235769-1r | | | 22,823,685 | 22,117,500 | 6.85 | 6.64 | 96.91 | 0.03 | 98.16 | 94.5 | 47.28 |
| | | S12-2_Root | FRAS202235767-1r | | | 20,355,828 | 19,761,200 | 6.11 | 5.93 | 97.08 | 0.03 | 98.08 | 94.3 | 45.86 |

| Morphs code | Voucher | Population tissues | Library | Pooled libraries | Read contents mapping rate (%) | Raw Reads | Clean Reads | Raw Base (G) | Clean Base (G) | Effective Rate (%) | Error Rate (%) | Q20 (%) | Q30 (%) | GC Content (%) |
|---|---|---|---|---|---|---|---|---|---|---|---|---|---|---|
| | | S12-2_Stem | FRAS202235768-1r | | | 19,706,866 | 19,062,308 | 5.91 | 5.72 | 96.73 | 0.03 | 97.99 | 94.17 | 46.64 |
| | | S12-3_Flower | FRAS202235774-1r | S12-3 merged | 95.95% | 19,926,078 | 19,241,324 | 5.98 | 5.77 | 96.56 | 0.03 | 97.97 | 94.14 | 47.63 |
| | | S12-3_Leaf | FRAS202235773-1r | | | 21,698,467 | 21,098,612 | 6.51 | 6.33 | 97.24 | 0.03 | 97.94 | 93.97 | 47.14 |
| | | S12-3_Root | FRAS202235771-1r | | | 21,242,360 | 20,509,189 | 6.37 | 6.15 | 96.55 | 0.03 | 97.9 | 93.89 | 45.8 |
| | | S12-3_Stem | FRAS202235772-1r | | | 22,462,819 | 21,794,291 | 6.74 | 6.54 | 97.02 | 0.03 | 98.04 | 94.25 | 46.05 |
| | | S12-4_Flower | FRAS202235778-1r | S12-4 merged | 96.00% | 19,583,648 | 18,881,638 | 5.88 | 5.66 | 96.42 | 0.03 | 98.06 | 94.28 | 46.9 |
| | | S12-4_Leaf | FRAS202235777-1r | | | 23,358,450 | 22,588,134 | 7.01 | 6.78 | 96.7 | 0.03 | 98.1 | 94.41 | 46.85 |
| | | S12-4_Root | FRAS202235775-1r | | | 19,794,108 | 19,163,465 | 5.94 | 5.75 | 96.81 | 0.03 | 98.12 | 94.42 | 46.1 |
| | | S12-4_Stem | FRAS202235776-1r | | | 21,750,481 | 21,156,098 | 6.53 | 6.35 | 97.27 | 0.03 | 97.94 | 94.01 | 46.59 |
| YY | Deng-9726 | S19-1_Flower | FRAS202235782-1r | S19-1 merged | 96.17% | 24,225,697 | 23,248,742 | 7.27 | 6.97 | 95.97 | 0.03 | 97.96 | 94.09 | 46.57 |
| | | S19-1_Leaf | FRAS202235781-1r | | | 23,641,987 | 22,914,757 | 7.09 | 6.87 | 96.92 | 0.03 | 98 | 94.11 | 47.03 |
| | | S19-1_Root | FRAS202235779-1r | | | 21,536,991 | 20,835,633 | 6.46 | 6.25 | 96.74 | 0.03 | 98.15 | 94.44 | 45.57 |
| | | S19-1_Stem | FRAS202235780-1r | | | 22,324,082 | 21,648,225 | 6.7 | 6.49 | 96.97 | 0.03 | 98.06 | 94.29 | 46.09 |
| | | S19-2_Flower | FRAS202235786-1r | S19-2 merged | 96.37% | 20,698,952 | 19,823,586 | 6.21 | 5.95 | 95.77 | 0.03 | 98.02 | 94.22 | 46.54 |
| | | S19-2_Leaf | FRAS202235785-1r | | | 23,276,629 | 22,381,621 | 6.98 | 6.71 | 96.15 | 0.03 | 98.1 | 94.38 | 46.54 |
| | | S19-2_Root | FRAS202235783-1r | | | 21,761,410 | 20,844,617 | 6.53 | 6.25 | 95.79 | 0.02 | 98.16 | 94.5 | 45.75 |
| | | S19-2_Stem | FRAS202235784-1r | | | 24,136,492 | 23,180,660 | 7.24 | 6.95 | 96.04 | 0.03 | 98.1 | 94.39 | 46.71 |

| Morphs code | Voucher | Population tissues | Library | Pooled libraries | Read contents mapping rate (%) | Raw Reads | Clean Reads | Raw Base (G) | Clean Base (G) | Effective Rate (%) | Error Rate (%) | Q20 (%) | Q30 (%) | GC Content (%) |
|---|---|---|---|---|---|---|---|---|---|---|---|---|---|---|
| | | S19-4_Flower | FRAS202235790-1r | S19-4 merged | 96.29% | 25,950,316 | 24,962,899 | 7.79 | 7.49 | 96.19 | 0.03 | 98.05 | 94.29 | 46.91 |
| | | S19-4_Leaf | FRAS202235789-1r | | | 21,295,363 | 20,593,185 | 6.39 | 6.18 | 96.7 | 0.03 | 98.09 | 94.39 | 46.92 |
| | | S19-4_Root | FRAS202235787-1r | | | 21,886,391 | 21,112,138 | 6.57 | 6.33 | 96.46 | 0.03 | 98.03 | 94.22 | 45.7 |
| | | S19-4_Stem | FRAS202235788-1b | | | 30,553,650 | 29,415,292 | 9.17 | 8.82 | 96.27 | 0.03 | 98.06 | 94.51 | 46.69 |

Note: Five different morphs: PP, Pure pink morph; RW, Red-white morph; RWh, Red-white transgressive mroph; YW, Yellow-white morph; YY, Pure yellow morph. Q20 – phred score that the base call accuracy of 99%; Q30 – phred score that the base call accuracy of 99.9%; biological replicates are written after the population code and before tissues name, for instance in S19-4_Stem, S19 is the population code of given voucher followed by 4 as its biological replicate code for the given tissue Stem.

**Supplementary Table S3** Statistics of the Trinity assembly of transcriptomes for four sample tissues of four different morphs of *Stellera chamaejasme*.

| Morphs code | Trinity Libraries | Counts of transcripts | | Stats based on ALL transcript contigs | | | | | | | | | Stats based on ONLY LONGEST ISOFORM per 'GENE' | | | | | | | |
|---|---|---|---|---|---|---|---|---|---|---|---|---|---|---|---|---|---|---|---|---|
| | | Total genes | Total transcripts | %GC | N10 | N20 | N30 | N40 | N50 | Median contig length (bp) | Average contig | Total assembled bases | N10 | N20 | N30 | N40 | N50 | Median contig length (bp) | Average contig | Total assembled bases |
| PP | PPRoot | 320625 | 560901 | 41.87 | 3879 | 2980 | 2420 | 1995 | 1637 | 519 | 931.68 | 522581568 | 3735 | 2809 | 2206 | 1731 | 1284 | 334 | 689.8 | 221166691 |
| | PPStem | 157537 | 352047 | 42.06 | 4300 | 3357 | 2750 | 2295 | 1909 | 694 | 1128.33 | 397226540 | 4382 | 3398 | 2742 | 2217 | 1712 | 415 | 853.82 | 134507457 |
| | PPLeaf | 181665 | 375563 | 42.11 | 4247 | 3284 | 2686 | 2238 | 1850 | 615 | 1060.2 | 398171986 | 4278 | 3269 | 2609 | 2081 | 1554 | 373 | 774.99 | 140788672 |
| | PPFlower | 239132 | 434300 | 42.83 | 4141 | 3220 | 2615 | 2163 | 1774 | 535 | 982.64 | 426762373 | 4069 | 3068 | 2406 | 1847 | 1300 | 345 | 701.03 | 167638893 |
| | PPmerged | 507384 | 899202 | 41.83 | 4181 | 3113 | 2456 | 1953 | 1537 | 452 | 860.6 | 773849740 | 3923 | 2783 | 2059 | 1498 | 1025 | 318 | 626.69 | 317973312 |
| RW | RWRoot | 186003 | 393406 | 42.36 | 4082 | 3180 | 2600 | 2160 | 1775 | 590 | 1017.81 | 400410770 | 4126 | 3175 | 2531 | 2011 | 1490 | 356 | 747.52 | 139040619 |
| | RWStem | 169508 | 385250 | 42.05 | 4332 | 3369 | 2750 | 2283 | 1885 | 653 | 1097.91 | 422969165 | 4434 | 3413 | 2737 | 2193 | 1664 | 404 | 830.51 | 140778130 |
| | RWLeaf | 165218 | 370114 | 42.05 | 4272 | 3317 | 2709 | 2256 | 1859 | 642 | 1083.1 | 400871823 | 4357 | 3345 | 2684 | 2152 | 1635 | 401 | 821.02 | 135646758 |
| | RWFlower | 188821 | 391890 | 42.41 | 4158 | 3242 | 2643 | 2195 | 1808 | 583 | 1027.58 | 402697631 | 4204 | 3233 | 2562 | 2036 | 1517 | 361 | 755.06 | 142570480 |
| | RWmerged | 296855 | 660173 | 41.66 | 4456 | 3360 | 2667 | 2137 | 1670 | 493 | 925.65 | 611091588 | 4449 | 3295 | 2500 | 1803 | 1178 | 334 | 677.29 | 201057221 |
| RWh | RWhRoot | 108825 | 208489 | 42.86 | 3878 | 3047 | 2538 | 2155 | 1834 | 756 | 1121.49 | 233818160 | 3990 | 3102 | 2556 | 2130 | 1748 | 387 | 846.6 | 92131075 |
| | RWhStem | 96266 | 184961 | 42.88 | 3579 | 2856 | 2405 | 2065 | 1777 | 813 | 1123.77 | 207853427 | 3699 | 2929 | 2450 | 2085 | 1743 | 420 | 879.83 | 84697359 |
| | RWhLeaf | 108487 | 217956 | 42.58 | 3661 | 2901 | 2435 | 2087 | 1785 | 798 | 1123.55 | 244884989 | 3809 | 2999 | 2495 | 2108 | 1741 | 413 | 865.64 | 93910830 |
| | RWhFlower | 140131 | 252981 | 42.9 | 3781 | 2972 | 2475 | 2103 | 1786 | 652 | 1051.12 | 265914055 | 3843 | 2978 | 2438 | 2024 | 1615 | 359 | 773.83 | 108437021 |

| **Morphs code** | **Trinity Libraries** | **Counts of transcripts** | | **Stats based on ALL transcript contigs** | | | | | | | | | **Stats based on ONLY LONGEST ISOFORM per 'GENE'** | | | | | | | |
|---|---|---|---|---|---|---|---|---|---|---|---|---|---|---|---|---|---|---|---|---|
| | | **Total genes** | **Total transcripts** | **% GC** | **N10** | **N20** | **N30** | **N40** | **N50** | **Median contig length (bp)** | **Average contig** | **Total assembled bases** | **N10** | **N20** | **N30** | **N40** | **N50** | **Median contig length (bp)** | **Average contig** | **Total assembled bases** |
| | RWhmerged | 189466 | 351859 | 42.35 | 4285 | 3345 | 2765 | 2327 | 1943 | 619 | 1086.98 | 382464553 | 4297 | 3299 | 2649 | 2125 | 1597 | 349 | 759.42 | 143883467 |
| YW | YWRoot | 257107 | 502670 | 42.48 | 4659 | 3617 | 2966 | 2458 | 2025 | 542 | 1065.36 | 535525117 | 4267 | 3213 | 2474 | 1854 | 1220 | 298 | 644.67 | 165750093 |
| | YWStem | 165003 | 415854 | 41.88 | 4955 | 3863 | 3189 | 2660 | 2219 | 792 | 1291.15 | 536931522 | 4706 | 3638 | 2917 | 2336 | 1799 | 417 | 872.6 | 143981363 |
| | YWLeaf | 160403 | 398129 | 41.89 | 4843 | 3767 | 3112 | 2602 | 2175 | 776 | 1265.51 | 503835283 | 4621 | 3553 | 2850 | 2291 | 1763 | 413 | 861.91 | 138252776 |
| | YWFlower | 194971 | 447961 | 42.03 | 4895 | 3807 | 3133 | 2614 | 2175 | 694 | 1212.68 | 543231867 | 4600 | 3520 | 2786 | 2188 | 1611 | 374 | 789.69 | 153966548 |
| | YWmerged | 370367 | 797966 | 41.78 | 5104 | 3899 | 3136 | 2543 | 2024 | 502 | 1029.77 | 821720173 | 4591 | 3309 | 2419 | 1639 | 1020 | 301 | 619.85 | 229573454 |
| YY | YYRoot | 205255 | 428982 | 42.1 | 4309 | 3329 | 2695 | 2219 | 1813 | 567 | 1016.58 | 436094639 | 4342 | 3302 | 2595 | 2025 | 1460 | 352 | 740.24 | 151937679 |
| | YYStem | 199960 | 474533 | 42.13 | 4544 | 3542 | 2892 | 2405 | 1984 | 634 | 1114.57 | 528899730 | 4364 | 3326 | 2629 | 2062 | 1495 | 365 | 758.19 | 151606901 |
| | YYLeaf | 180981 | 441075 | 42.12 | 4544 | 3528 | 2886 | 2398 | 1980 | 646 | 1125.13 | 496267473 | 4376 | 3354 | 2659 | 2094 | 1559 | 382 | 787.26 | 142479375 |
| | YYFlower | 277876 | 550323 | 43.31 | 4322 | 3337 | 2714 | 2246 | 1838 | 556 | 1018.14 | 560306650 | 4042 | 3015 | 2340 | 1783 | 1244 | 353 | 698.71 | 194155635 |
| | YYmerged | 427096 | 936698 | 42.26 | 4806 | 3628 | 2889 | 2305 | 1807 | 484 | 955.16 | 894693730 | 4317 | 3089 | 2243 | 1540 | 1001 | 322 | 631.77 | 269826783 |

N50 – length of the smallest contig in the set that contains the fewest (largest) contigs whose combined length represents at least 50% of the assembly and so on for N10, N20, N30, N40.

**Supplementary Table S4** Statistics of the filtering of transcriptomes/pan- transcriptomes for four sample tissues of four different morphs of *Stellera chamaejasme*.

| Morphs code | Trinity Libraries | Counts of transcripts | | | Stats based on ALL transcript contigs | | | | | | | | Stats based on ONLY LONGEST ISOFORM per 'GENE' | | | | | | | |
|---|---|---|---|---|---|---|---|---|---|---|---|---|---|---|---|---|---|---|---|---|
| | | Total genes | Total transcripts | %GC | N10 | N20 | N30 | N40 | N50 | Median contig length (bp) | Average contig (bp) | Total assembled bases | N10 | N20 | N30 | N40 | N50 | Median contig length (bp) | Average contig (bp) | Total assembled bases |
| PP | PPRoot | 274790 | 394065 | 41.78 | 3884 | 2979 | 2413 | 1980 | 1606 | 430 | 859.63 | 338748212 | 3821 | 2876 | 2260 | 1774 | 1307 | 324 | 684.63 | 188128207 |
| | PPStem | 130759 | 224291 | 41.86 | 4400 | 3462 | 2858 | 2394 | 2000 | 628 | 1115.64 | 250228050 | 4511 | 3519 | 2862 | 2338 | 1849 | 416 | 887.55 | 116055351 |
| | PPLeaf | 152894 | 246683 | 42.02 | 4325 | 3365 | 2764 | 2311 | 1915 | 535 | 1027.63 | 253498848 | 4393 | 3372 | 2709 | 2188 | 1679 | 370 | 796.53 | 121785314 |
| | PPFlower | 208133 | 301374 | 42.72 | 4175 | 3253 | 2640 | 2177 | 1771 | 440 | 908.44 | 273781633 | 4137 | 3149 | 2473 | 1909 | 1350 | 336 | 699.39 | 145565622 |
| | PPmerged | 435465 | 621338 | 41.93 | 4272 | 3196 | 2520 | 1998 | 1556 | 389 | 810.9 | 503840573 | 4062 | 2902 | 2155 | 1564 | 1053 | 311 | 626.46 | 272800228 |
| RW | RWRoot | 154808 | 252129 | 42.21 | 4135 | 3246 | 2664 | 2220 | 1831 | 510 | 980.32 | 247166481 | 4209 | 3273 | 2631 | 2112 | 1612 | 357 | 770.61 | 119296269 |
| | RWStem | 141363 | 243016 | 41.85 | 4450 | 3484 | 2856 | 2386 | 1984 | 590 | 1086.37 | 264005365 | 4543 | 3524 | 2840 | 2311 | 1801 | 406 | 862.19 | 121881176 |
| | RWLeaf | 137758 | 235320 | 41.88 | 4364 | 3420 | 2808 | 2350 | 1951 | 580 | 1068.68 | 251481863 | 4457 | 3440 | 2783 | 2265 | 1760 | 403 | 851.34 | 117279454 |
| | RWFlower | 160706 | 256483 | 42.19 | 4242 | 3317 | 2718 | 2251 | 1851 | 495 | 979.54 | 251234335 | 4296 | 3312 | 2648 | 2122 | 1614 | 358 | 770.23 | 123780950 |
| | RWmerged | 245652 | 411152 | 41.7 | 4665 | 3558 | 2855 | 2320 | 1843 | 448 | 931.97 | 383181942 | 4643 | 3462 | 2676 | 2007 | 1352 | 336 | 707.92 | 173901466 |
| RWh | RWhRoot | 91618 | 144293 | 42.69 | 3880 | 3064 | 2555 | 2172 | 1844 | 641 | 1064.47 | 153596189 | 4037 | 3154 | 2609 | 2186 | 1801 | 382 | 858.79 | 78680947 |
| | RWhStem | 80840 | 129150 | 42.64 | 3635 | 2897 | 2435 | 2092 | 1795 | 711 | 1077.74 | 139190660 | 3774 | 3000 | 2514 | 2135 | 1792 | 421 | 896.01 | 72433409 |
| | RWhLeaf | 90172 | 148930 | 42.37 | 3748 | 2977 | 2495 | 2142 | 1832 | 717 | 1097.93 | 163515190 | 3885 | 3073 | 2568 | 2178 | 1811 | 419 | 896.54 | 80843177 |
| | RWhFlower | 120419 | 179485 | 42.7 | 3822 | 3004 | 2500 | 2119 | 1792 | 527 | 983.82 | 176580054 | 3908 | 3042 | 2492 | 2067 | 1657 | 353 | 777.92 | 93676458 |

| **Morphs code** | **Trinity Libraries** | **Counts of transcripts** | | | **Stats based on ALL transcript contigs** | | | | | | | | **Stats based on ONLY LONGEST ISOFORM per 'GENE'** | | | | | | | |
|---|---|---|---|---|---|---|---|---|---|---|---|---|---|---|---|---|---|---|---|
| | | **Total genes** | **Total transcripts** | **%GC** | **N10** | **N20** | **N30** | **N40** | **N50** | **Median contig length (bp)** | **Average contig (bp)** | **Total assembled bases** | **N10** | **N20** | **N30** | **N40** | **N50** | **Median contig length (bp)** | **Average contig (bp)** | **Total assembled bases** |
| | RWhmerged | 162871 | 249796 | 42.29 | 4339 | 3406 | 2816 | 2372 | 1986 | 510 | 1032.92 | 258019754 | 4378 | 3389 | 2744 | 2237 | 1715 | 338 | 769.46 | 125322754 |
| YW | YWRoot | 215419 | 325293 | 42.66 | 4678 | 3640 | 2991 | 2478 | 2037 | 414 | 973.9 | 316804002 | 4394 | 3342 | 2608 | 1994 | 1362 | 287 | 653.35 | 140743494 |
| | YWStem | 128165 | 239669 | 41.72 | 4999 | 3952 | 3284 | 2758 | 2319 | 733 | 1287.46 | 308563273 | 4870 | 3804 | 3088 | 2513 | 2008 | 434 | 940.76 | 120572972 |
| | YWLeaf | 125059 | 231663 | 41.75 | 4881 | 3841 | 3189 | 2686 | 2256 | 717 | 1255.47 | 290845511 | 4783 | 3726 | 3022 | 2468 | 1958 | 429 | 925.81 | 115780692 |
| | YWFlower | 154915 | 267996 | 41.9 | 4954 | 3889 | 3216 | 2694 | 2251 | 602 | 1178.56 | 315850124 | 4771 | 3687 | 2952 | 2375 | 1815 | 378 | 835.28 | 129397093 |
| | YWmerged | 301861 | 482992 | 42.03 | 5243 | 4044 | 3282 | 2685 | 2164 | 418 | 991.79 | 479027894 | 4793 | 3534 | 2664 | 1892 | 1182 | 296 | 641.77 | 193723876 |
| YY | YYRoot | 172925 | 279313 | 42.06 | 4423 | 3441 | 2791 | 2317 | 1903 | 494 | 990.57 | 276679774 | 4466 | 3427 | 2712 | 2157 | 1601 | 347 | 760.42 | 131496313 |
| | YYStem | 161408 | 275756 | 41.91 | 4576 | 3604 | 2949 | 2463 | 2038 | 540 | 1066.69 | 294145128 | 4498 | 3443 | 2756 | 2200 | 1650 | 369 | 791.06 | 127683193 |
| | YYLeaf | 144201 | 252839 | 41.89 | 4632 | 3613 | 2965 | 2471 | 2050 | 568 | 1093.31 | 276430858 | 4518 | 3479 | 2794 | 2230 | 1710 | 392 | 828.57 | 119481090 |
| | YYFlower | 224539 | 335379 | 43.1 | 4323 | 3337 | 2711 | 2232 | 1806 | 457 | 930.13 | 311945201 | 4130 | 3106 | 2437 | 1869 | 1328 | 351 | 712.5 | 159984414 |
| | YYmerged | 342094 | 544648 | 42.33 | 4968 | 3791 | 3033 | 2439 | 1921 | 421 | 924.16 | 503343972 | 4535 | 3290 | 2443 | 1725 | 1115 | 321 | 654.56 | 223920443 |

N50 – length of the smallest contig in the set that contains the fewest (largest) contigs whose combined length represents at least 50% of the assembly and so on for N10, N20, N30, N40.

**Supplementary Table S5** OrthoFinder summary statistics for transcriptome assemblies across floral morphs of *Stellera chamaejasme.*

| **Metrics** | **Pure pink morph (PP)** | **Red-white transgressive morph (RWh)** | **Red-white morph (RW)** | **Yellow-white morph (YW)** | **Pure yellow morph (YY)** |
|---|---|---|---|---|---|
| Total number of genes | 243,920 | 125,060 | 164,898 | 215,661 | 234,140 |
| Genes assigned to orthogroups | 220,437 | 117,546 | 154,133 | 197,163 | 211,523 |
| Unassigned genes | 23,483 | 7,514 | 10,765 | 18,498 | 22,617 |
| % genes in orthogroups | 90.4 | 94 | 93.5 | 91.4 | 90.3 |
| % unassigned genes | 9.6 | 6 | 6.5 | 8.6 | 9.7 |
| Orthogroups containing species | 106,852 | 71,751 | 86,763 | 96,830 | 106,711 |
| % orthogroups containing species | 66.5 | 44.7 | 54 | 60.3 | 66.4 |
| Species-specific orthogroups | 9,759 | 1,818 | 3,220 | 5,323 | 7,078 |
| Genes in species-specific orthogroups | 38,815 | 4,614 | 8,412 | 15,176 | 19,457 |
| % genes in species-specific orthogroups | 15.9 | 3.7 | 5.1 | 7 | 8.3 |

**Supplementary Table S6** Representative morph-associated DEGs and their functional roles in pigment biosynthesis and regulatory processes.

| Morph | Gene (abbrev.) | Full name | Functional role | Pathway / Process | Key references |
|---|---|---|---|---|---|
| **PP** | *10HGO* | 10-hydroxygeraniol oxidoreductase | Secondary metabolite synthesis | Terpenoid/secondary metabolism | 1 |
| | *BGLU44* | β-Glucosidase 44 | Secondary metabolite activation | Secondary metabolism | 2 |
| | *CAD* | Cinnamyl alcohol dehydrogenase | Lignin biosynthesis | Phenylpropanoid pathway | 3 |
| | *HIUH* | Hydantoinase-like protein | Nitrogen metabolism | Stress/metabolism | 4 |
| | *NCED1* | 9-cis-epoxycarotenoid dioxygenase | ABA biosynthesis | Hormone signaling | 5 |
| | *PER31* | Peroxidase 31 | Redox regulation | Stress response | 6 |
| | *PER52* | Peroxidase 52 | ROS detoxification | Oxidative stress | 6 |
| | *PER65* | Peroxidase 65 | Redox regulation | Stress response | 6 |
| | *pod* | Peroxidase (POD) | Redox regulation | Stress response | 6 |
| | *4CL2* | 4-Coumarate-CoA ligase 2 | Activates hydroxycinnamic acids | Phenylpropanoid metabolism | 7 |
| | *AN3* | Anthocyanin regulator 3 | Pigment regulation | Flavonoid regulation | 8 |
| | *CCR1* | Cinnamoyl-CoA reductase 1 | Controls lignin/phenylpropanoid flux | Phenylpropanoid metabolism | 9 |
| | *F3H* | Flavanone 3-hydroxylase | Converts flavanones to dihydroflavonols | Anthocyanin biosynthesis | 10 |
| | *HCT* | Hydroxycinnamoyl transferase | Intermediate formation in lignin/flavonoid pathways | Phenylpropanoid metabolism | 11 |
| | *HST* | Hydroxycinnamoyl-CoA shikimate transferase-like | Phenylpropanoid flux regulation | Phenylpropanoid metabolism | 11 |
| | *L1AT* | Anthocyanin acyltransferase | Anthocyanin stabilization/modification | Flavonoid modification | 8 |
| | *PER12* | Peroxidase 12 | ROS detoxification | Oxidative stress response | 6 |
| | *PRDX6* | Peroxiredoxin 6 | Redox homeostasis | Antioxidant defense | 12 |

| Morph | Gene (abbrev.) | Full name | Functional role | Pathway / Process | Key references |
|---|---|---|---|---|---|
| | *WRKY6* | WRKY transcription factor 6 | Stress and secondary metabolism regulation | Transcriptional regulation | 13 |
| **RW** | *4CL1* | 4-Coumarate-CoA ligase 1 | Phenylpropanoid activation | Phenylpropanoid metabolism | 7 |
| | *hflX* | Ribosome-associated GTPase | Stress adaptation | Translation/stress | 14 |
| | *WRKY6* | WRKY transcription factor 6 | Stress/transcription regulation | Regulatory signaling | 13 |
| | *BGLU18* | β-Glucosidase 18 | Hydrolysis of secondary metabolite conjugates | Secondary metabolism | 2 |
| | *BGLU24* | β-Glucosidase 24 | Secondary metabolite activation | Secondary metabolism | 2 |
| | *CCD8* | Carotenoid cleavage dioxygenase 8 | Carotenoid degradation/cleavage | Carotenoid metabolism | 15 |
| | *COMT* | Caffeic acid O-methyltransferase | Methylation of lignin/flavonoid intermediates | Phenylpropanoid metabolism | 9 |
| | *CTOMT1* | Caffeoyl-CoA O-methyltransferase 1 | Lignin/flavonoid methylation | Phenylpropanoid metabolism | 7 |
| | *GT5* | UDP-glycosyltransferase family protein | Glycosylation of pigments/metabolites | Secondary metabolite modification | 16 |
| | *HRPN* | Hypersensitive response protein-like | Defense/stress signaling | Stress response | 17 |
| **RWh** | *BETA-OHASE* | β-Carotene hydroxylase | Conversion of β-carotene to xanthophylls | Carotenoid metabolism | 18 |
| | *L21AT* | Acyltransferase-like protein | Secondary metabolite modification | Flavonoid metabolism | 8 |
| | *UGT73C3* | UDP-glycosyltransferase 73C3 | Glycosylation of secondary metabolites | Secondary metabolism | 16 |
| | *BGLU31* | β-Glucosidase 31 | Secondary metabolite activation | Secondary metabolism | 2 |
| | *FGT* | Flavonoid | Flavonoid modification | Flavonoid | 16 |

| Morph | Gene (abbrev.) | Full name | Functional role | Pathway / Process | Key references |
|---|---|---|---|---|---|
| | | glycosyltransferase | | metabolism | |
| | *PER10* | Peroxidase 10 | Oxidative stress response | Redox metabolism | 6 |
| | *SKIP1* | SNW/SKI-interacting protein 1 | Transcription/splicing co-regulator | Stress and transcription regulation | 19 |
| | *TIFY6B* | TIFY domain protein 6B | Jasmonate signaling regulator | Hormone/stress signaling | 20 |
| **YW** | *PER11* | Peroxidase 11 | ROS detoxification | Stress response | 6 |
| | *PSY* | Phytoene synthase | Rate-limiting carotenoid synthesis | Carotenoid biosynthesis | 15 |
| | *TOP1B* | DNA topoisomerase I beta | DNA topology regulation | Transcription/replication | 21 |
| | *BBX22* | B-box zinc finger protein 22 | Light-regulated transcription | Light signaling/pigment regulation | 22 |
| | *CCOAOMT2* | Caffeoyl-CoA O-methyltransferase 2 | Phenylpropanoid methylation | Phenylpropanoid metabolism | 7 |
| | *F3GGT1* | Flavonoid 3-O-glucoside glucosyltransferase | Flavonoid glycosylation | Flavonoid modification | 16 |
| | *Z-ISO* | ζ-Carotene isomerase | Carotenoid biosynthesis isomerization step | Carotenoid biosynthesis | 23 |
| **YY** | *At1g65750* | Unknown Arabidopsis homolog | Unknown/putative enzyme | Unknown | 24 |
| | *CCD4* | Carotenoid cleavage dioxygenase 4 | Carotenoid degradation | Carotenoid metabolism | 15 |
| | *RG* | RG-like protein | Secondary metabolism | Unknown | 1 |
| | *BGLU5* | β-Glucosidase 5 | Secondary metabolite turnover | Secondary metabolism | 2 |
| | *BGLU6* | β-Glucosidase 6 | Secondary metabolite turnover | Secondary metabolism | 2 |
| | *CCOMT* | Caffeoyl-CoA O- | Phenylpropanoid metabolism | Phenylpropanoid | 7 |

| Morph | Gene (abbrev.) | Full name | Functional role | Pathway / Process | Key references |
|---|---|---|---|---|---|
| | | methyltransferase | | pathway | |
| | *FOX2* | Multifunctional β-oxidation protein | Fatty acid degradation/peroxisomal metabolism | Lipid metabolism | 25 |
| | *PER47* | Peroxidase 47 | ROS detoxification | Oxidative stress response | 6 |


**References**:

1. Dewick, P. M. *Medicinal natural products: A biosynthetic approach*. John Wiley & Sons (2002).
2. Xu, Z. et al. Functional genomic analysis of *Arabidopsis thaliana* glycoside hydrolase family 1. *Plant Mol. Biol.* **55**, 343–367 (2004).
3. Boerjan, W., Ralph, J. & Baucher, M. Lignin biosynthesis. *Annu. Rev. Plant Biol.* **54**, 519–546 (2003).
4. Hu, J. et al. Plant peroxisomes: biogenesis and function. *Plant Cell* **24**, 2279–2303 (2012).
5. Nambara, E. & Marion-Poll, A. Abscisic acid biosynthesis and catabolism. *Annu. Rev. Plant Biol.* **56**, 165–185 (2005).
6. Passardi, F., Cosio, C., Penel, C. & Dunand, C. Peroxidases have more functions than a Swiss army knife. *Plant Cell Rep.* **24**, 255–265 (2005).
7. Ehlting, J. et al. Three 4-coumarate: coenzyme A ligases in *Arabidopsis thaliana* represent two evolutionarily divergent classes in angiosperms. *Plant J.* **19**, 9–20 (1999).
8. Grotewold, E. The genetics and biochemistry of floral pigments. *Annu. Rev. Plant Biol.* **57**, 761–780 (2006).
9. Goujon, T. et al. Genes involved in the biosynthesis of lignin precursors in *Arabidopsis thaliana*. *Plant Physiol. Biochem.* **41**, 677–687 (2003).
10. Holton, T. A. & Cornish, E. C. Genetics and biochemistry of anthocyanin biosynthesis. *Plant Cell* **7**, 1071 (1995).
11. Pu, G., Zhou, B. & Xiang, F. Isolation and functional characterization of a *Lonicera japonica* hydroxycinnamoyl transferase involved in chlorogenic acid synthesis. *Biologia* **72**, 608–618 (2017).
12. Dietz, K. J. Peroxiredoxins in plants and cyanobacteria. *Antioxid. Redox Signal.* **15**, 1129–1159 (2011).
13. Rushton, P. J. et al. *WRKY* transcription factors. *Trends Plant Sci.* **15**, 247–258 (2010).
14. Zhang, Y. et al. *HflX* is a ribosome-splitting factor rescuing stalled ribosomes under stress conditions. *Nat. Struct. Mol. Biol.* **22**, 906–913 (2015).
15. Nisar, N. et al. Carotenoid metabolism in plants. *Mol. Plant* **8**, 68–82 (2015).
16. Bowles, D. et al. Glycosyltransferases: managers of small molecules. *Curr. Opin. Plant Biol*. **8**, 254–263 (2005).
17. Van Loon, L. C., Rep, M. & Pieterse, C. M. Significance of inducible defense-related proteins in infected plants. *Annu. Rev. Phytopathol.* **44**, 135–162 (2006).

18. Kim, J. et al. The evolution and function of carotenoid hydroxylases in *Arabidopsis*. *Plant Cell Physiol.* **50**, 463–479 (2009).
19. Cao, Y. & Ma, L. To splice or to transcribe: SKIP-mediated environmental fitness and development in plants. *Front. Plant Sci.* **10**, 1222 (2019).
20. Vanholme, B. et al. The *tify* family previously known as *ZIM*. *Trends Plant Sci*. **12**, 239–244 (2007).
21. Wang, J. C. Cellular roles of DNA topoisomerases: a molecular perspective. *Nat. Rev. Mol. Cell Biol.* **3**, 430–440 (2002).
22. Gangappa, S. N. & Botto, J. F. The BBX family of plant transcription factors. *Trends Plant Sci*. **19**, 460–470 (2014).
23. Chen, Y. et al. Isolation and characterization of the *Z-ISO* gene encoding a missing component of carotenoid biosynthesis in plants. *Plant Physiol*. **153**, 66–79 (2010).
24. Berardini, T. Z. et al. The *Arabidopsis* information resource: making and mining the "gold standard" annotated reference plant genome. *Genesis* **53**, 474–485 (2015).
25. Graham, I. A. Seed storage oil mobilization. *Annu. Rev. Plant Biol*. **59**, 115–142 (2008).

**Supplementary Table S7** Association between hybrid inheritance mode and variance inflation across transcriptional modules.

| Mode | N | Median | Q1 | Q3 |
|---|---|---|---|---|
| Additive | 14 | 0.88 | 0.393 | 1.18 |
| Transgressive | 20 | 0.669 | 0.453 | 1.23 |
| Dominant_PP | 15 | 0.342 | 0.261 | 0.797 |
| Dominant_RW | 4 | 0.34 | 0.292 | 0.777 |

Median and interquartile range (IQR) of hybrid variance inflation indices are shown for transcriptional modules grouped by inheritance mode. N indicates the number of co-expression modules assigned to each inheritance category. Inflation indices were calculated as the ratio of within-population variance of the transgressive morph (RWh) to the mean parental variance across non-transgressive morphotypes. Differences between transgressive and non-transgressive modules were evaluated using a Wilcoxon rank-sum test ($W = 267$, $p = 0.254$).
